# Light-induced Patterning of Electroactive Bacterial Biofilms

**DOI:** 10.1101/2021.12.20.473588

**Authors:** Fengjie Zhao, Marko S. Chavez, Kyle L. Naughton, Christina M. Cole, Jeffrey A. Gralnick, Mohamed Y. El-Naggar, James Q. Boedicker

## Abstract

Electroactive bacterial biofilms can function as living biomaterials that merge the functionality of living cells with electronic components. However, the development of such advanced living electronics has been challenged by the inability to control the geometry of electroactive biofilms relative to solid-state electrodes. Here, we developed a lithographic strategy to pattern conductive biofilms of *Shewanella oneidensis* by controlling aggregation protein CdrAB expression with a blue light-induced genetic circuit. This controlled deposition enabled *S. oneidensis* biofilm patterning on transparent electrode surfaces and measurements demonstrated tunable biofilm conduction dependent on pattern size. Controlling biofilm geometry also enabled us, for the first time, to quantify the intrinsic conductivity of living *S. oneidensis* biofilms and experimentally confirm predictions based on simulations of a recently proposed collision-exchange electron transport mechanism. Overall, we developed a facile technique for controlling electroactive biofilm formation on electrodes, with implications for both studying and harnessing bioelectronics.

## Introduction

Exoelectrogens can use extracellular electron transfer (EET) pathways to gain energy by respiring solid surfaces such as minerals and electrodes in anoxic environments^1, 2^. *Shewanella oneidensis* MR-1 is one of the most studied model exoelectrogenic organisms, whose EET pathway includes a network of multiheme *c*-type cytochromes to route electrons from the cellular interior to external surfaces^3–9^. Electrons are transferred to external, solid electron acceptors indirectly via soluble flavins that serve as electron shuttles and directly via contact with the cell surface cytochromes, flavin-cytochrome complexes, or along membrane extensions that are proposed to act as nanowires^10–13^.

When performing direct electron transfer, exoelectrogens are able to colonize electrode surfaces and form living conductive biofilms, which allow for long-distance electron transport across neighboring cells on the micrometer-scale^14^. Electrochemical techniques have proved valuable for investigating the electron transport mechanisms in these biofilms, demonstrating that redox gradient-driven electron hopping is the basis of biofilm conduction in both *Geobacter* and *Shewanella*^15–17^. Such electroactive biofilms, acting as living biomaterials, have found applications in microbial fuel cells, microbial electrosynthesis cells, nanoparticle templating, and bioremediation^18, 19^. In addition to these sustainability applications, electroactive biofilms have recently been proposed to form the basis of living electronics, hybrid devices that integrate cells on electrodes to combine the functionality of a biological system (e.g. for biosensing, biocomputing, or data storage) with the readout, stimulation, and amplification advantages of electronic circuitry^14, 20, 21^. To realize such hybrid technologies, it is important to develop new tools for controlling electroactive biofilm formation in relation to solid-state electrodes with high spatial resolution.

Synthetic biology strategies have recently been developed for regulating the EET capabilities of *S. oneidensis*, such as tuning *mtrCAB* expression level with a native inducible system^22^ and using clustered regularly interspaced short palindromic repeats interference (CRISPRi) to regulate the transcription of *mtrA*^23^. In a recent study, a quorum sensing-based population-state decision (PSD) system was constructed to intelligently reprogram the EET network^24^. The EET pathway has also been reconstructed successfully in *Escherichia coli* through heterologous expression of the MtrCAB complex^25^. The EET ability of engineered *E. coli* can be further enhanced through coexpression of CymA, adding exogenous flavins or modifying the cytochrome *c* maturation pathway^26, 27^. Although much progress has been made in manipulating the EET machinery of *Shewanella*, very little work has been done in tuning EET through the controlled formation of conductive biofilms on solid-state electrodes. Many downstream applications and fundamental experimental measurements would benefit from the ability to precisely pattern electroactive biofilms on electrode surfaces.

In recent years, optogenetic approaches have been developed for spatial control of gene expression in bacteria as a response to illumination with light^28–30^. These techniques provide possible strategies to dynamically control cell-cell adhesions for patterning bacterial biofilms with high spatial and temporal precision. Some studies have utilized light-induced genetic circuits to control the expression of aggregation proteins that mediate cell-cell adhesions and realize bacterial biofilm patterning. A technique called “Biofilm Lithography” was recently developed, which used the blue light-induced pDawn genetic circuit to control the expression of *E. coli* cellsurface adhesin Ag43 in order to pattern *E. coli* biofilms with high spatial resolution^31^. Biofilm patterning of *E. coli* with high spatial resolution on various types of material surfaces has also been achieved by controlling the biosynthesis of curli fiber in response to illumination with various wavelengths of light^32^. Optogenetic tools can also directly control protein activities to adjust bacterial biofilm formation, such as activating proteins involved in cyclic dimeric guanosine monophosphate (*c*-di-GMP) synthesis or hydrolysis^33^ and toggling cell-cell adhesion through photo-switchable protein heterodimers^34^. Although the ability to pattern *E. coli* biofilms exists, electrochemical measurements of engineered, EET capable *E. coli* have yielded low electrical current^25^ and had limitations for further applications compared with naturally electroactive bacteria. Similarly, biofilm patterning techniques have not previously been adapted for naturally electroactive bacteria. Inspired by these robust optogenetic patterning techniques, we set out to implement a similar strategy in the EET model organism *S. oneidensis* to enable light-patterning of naturally electroactive biofilms. Thus, biofilm size-dependent tunability of electron transfer in conductive biofilms and the accurate determination of biofilm conductivity from the defined patterns can be possible.

In this work, we introduced the blue light-induced pDawn genetic circuit into *S. oneidensis* to control the expression of aggregation proteins CdrAB. CdrAB is a set of proteins found in *Pseudomonas aeruginosa* that helps to regulate adhesion and biofilm formation^35, 36^. Biofilm patterning of the resulting CdrAB strain was first achieved on both plastic and glass substrates under blue light illumination. Next, we confirmed that it was possible to pattern the CdrAB strain onto transparent, indium tin oxide (ITO) electrode surfaces and that these patterned biofilms remained electrochemically active. By patterning biofilms on ITO interdigitated array (IDA) microelectrodes and performing electrochemical gating measurements, we demonstrated that biofilm conduction could be tuned by controlling the biofilm dimensions. Finally, this precise control of biofilm pattern geometry allowed us to determine the intrinsic redox conductivity of the patterned biofilms. To our knowledge, this is the first demonstration of light-based lithography of naturally electroactive biofilms on electrodes.

## Results

### The pDawn genetic circuit enables blue light-induced protein expression in *S. oneidensis*

The light responsive pDawn genetic circuit has been used to control gene expression in *E. coli* and *P. aeruginosa*^29, 37^, but has yet to be tested in *S. oneidensis*. To verify whether the blue light-induced pDawn genetic circuit is functional in *S. oneidensis*, we expressed the red fluorescent protein mCherry with the pDawn plasmid (Fig. 1A) by irradiating the cells with blue light LEDs while culturing them in a shaking incubator. The relative fluorescence intensity of the mCherry strain was increased 58-fold under blue light illumination compared with that of those cells cultured in the dark (Fig. 1B). The mCherry strain grown in the dark and the untransformed wild type cells grown either in the dark or under blue light illumination exhibited very weak fluorescence intensity (Fig. 1B). Bright red fluorescence could be seen in the fluorescence microscopy images of mCherry cells grown under blue light illumination, while fluorescence was not detected from microscopy for the cells grown in the dark (Fig. 1C). Taken collectively, these results indicate that the pDawn light-activated transcriptional promoter can be used to optically regulate gene expression in *S. oneidensis*.

**Fig. 1.**
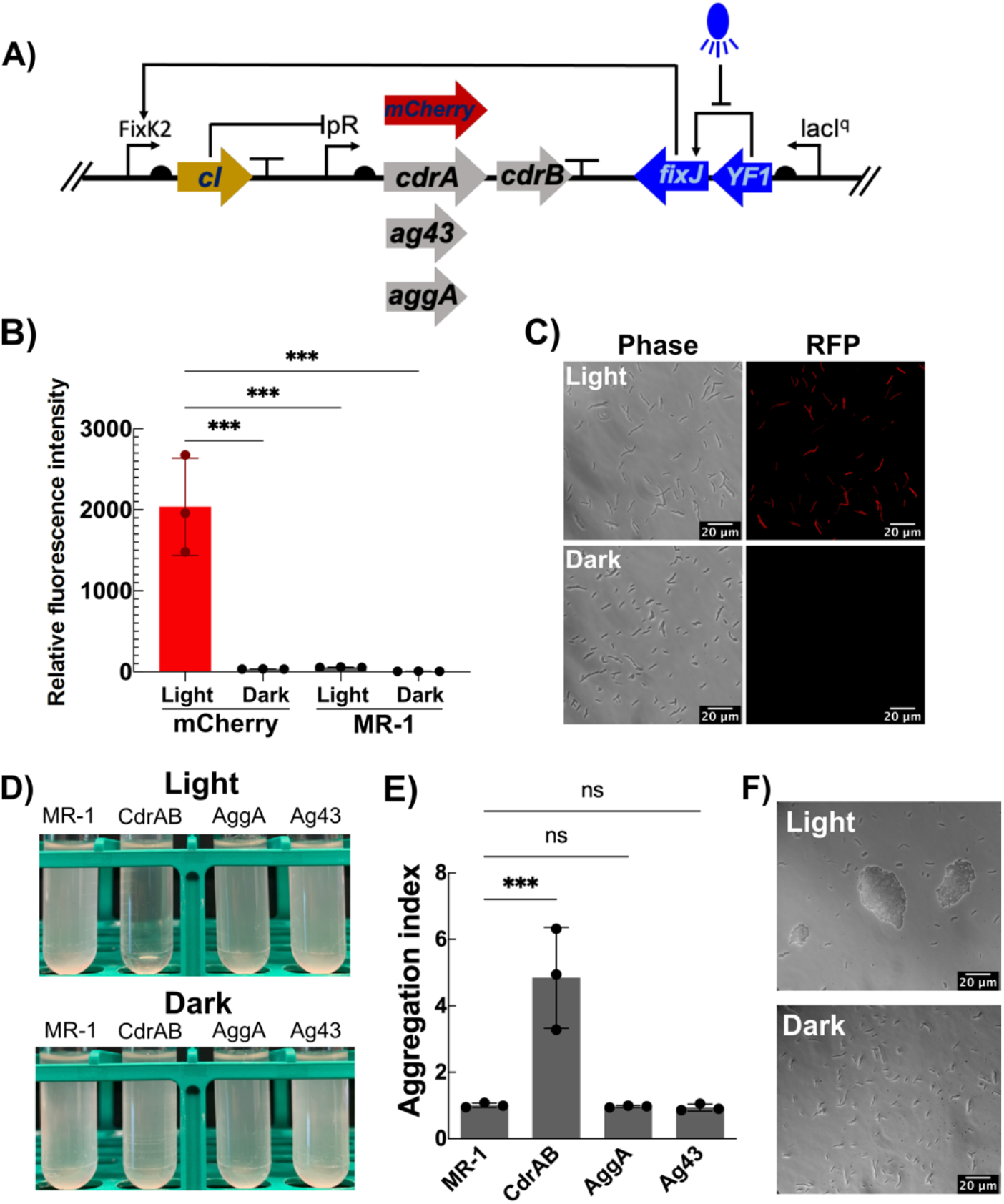
Characterization of the pDawn genetic circuit in *Shewanella* and selection of adhesive proteins for clumping *Shewanella* cells under blue light illumination. **A** The genetic circuit pDawn was used for light-induced expression of mCherry or different adhesive proteins. **B** Fluorescence intensity measurements of mCherry and wild type strains cultured under blue light and dark conditions, respectively. n = 3 independent biological experiments per group. *p* = 0.0001 for mCherry Light vs mCherry Dark, mCherry Light vs MR-1 Light and mCherry Light vs MR-1 Dark (one-way ANOVA with Dunnett’s multiple comparisons test). **C** Microscope observation for mCherry cells grown under blue light and dark conditions, respectively. **D** Cell clumping assay of *Shewanella* strains with different aggregation proteins expressed under blue light. **E** Aggregation indexes of different *Shewanella* strains. n = 3 independent biological experiments per group. *p* = 0.0007 for MR-1 vs CdrAB, *p* > 0.9999 for MR-1 vs AggA and MR-1 vs Ag43 (one-way ANOVA with Dunnett’s multiple comparisons test). **F** Microscope observation for cell clusters of the CdrAB strain cultured under blue light and dark conditions, respectively. Data are shown as mean ± SD. Significance is indicated as \*\*\**p* < 0.001 and ns (not significant) *p* > 0.05.

### The expression of CdrAB proteins can mediate cell-cell aggregations in *S. oneidensis*

One of the strategies for mediating bacterial cell-cell aggregations is to express outer membrane adhesive proteins involved in biofilm formation. Thus, several putative proteins that promote bacterial auto-aggregation and biofilm formation from different species were selected to be expressed in *S. oneidensis* under the control of the pDawn genetic circuit, including Ag43 from *E. coli*^38, 39^ and CdrAB from *P. aeruginosa* PAO1^35, 36^ (Fig. 1A). The native AggA aggregation protein from *S. oneidensis*^40, 41^ was also overexpressed using the pDawn genetic circuit (Fig. 1A).

Cells that cluster when grown in liquid culture will sediment to the bottom of culture tubes if not shaken, reducing the turbidity of the culture at the top of the tube. As shown in Fig. 1D, expression of CdrAB under blue light illumination triggered obvious cell-cell aggregation and sedimentation in *S. oneidensis*. However, we were not able to detect cell-cell aggregations when inducing the expression of Ag43 or the overexpression of AggA (Fig. 1D). The extent to which cell adhesion increased upon light-induced expression of the potential aggregation genes was quantified using an “aggregation index”. This aggregation index measures the reduction of the culture turbidity at the top of the culture tube after settling (see Methods). The aggregation index of 4.85, measured for the CdrAB strain, indicated significant cell-cell adhesion (Fig. 1E). However, the wild type host and strains with inducible expression of Ag43 and AggA showed an aggregation index near 1, indicating no increased sedimentation (Fig. 1E). Microscopy confirmed many large cell clusters were formed due to the cell-cell aggregations of the CdrAB strain grown under blue light illumination, while no large cell clusters were observed under the dark culturing condition (Fig. 1F). These results indicate that the expression of CdrAB proteins can be controlled by light and mediate cell-cell aggregations in *S. oneidensis*.

### Biofilm formation can be enhanced and patterned by expressing CdrAB in *S. oneidensis* under blue light

Bacterial biofilm formation is closely coupled to cell aggregation^34^. To verify that the biofilm formation of the CdrAB strain can be significantly enhanced when grown under blue light illumination, we performed a biofilm formation assay. Cells were grown in 96-well plates and exposed to blue light from a portable LED projector (Fig. 2A). Crystal violet staining was used to quantify biofilm formation^42^. The results showed that the CdrAB strain formed robust biofilms under blue light illumination, while weak biofilms were formed in the dark (Fig. 2B). Wild-type cells failed to form robust biofilms regardless of dark or blue light illuminated growth conditions (Fig. 2B). Next, we optimized biofilm formation by systematically testing the impact of illumination intensity and time. We found that the extent of CdrAB biofilm formation reached a maximum with 110 μW/cm^2^ light intensity and then decreased for light intensities exceeding 120 μW/cm^2^ when grown overnight for 16 h (Supplementary Fig. 1A). This may be due to higher levels of light intensity negatively impacting cellular growth, reducing biofilm formation despite higher expression of CdrAB. CdrAB biofilm formation peaked after about 16 h and then decreased with increasing incubation time when illuminated at 110 μW/cm^2^ (Supplementary Fig. 1B), indicating that biofilm formation changes dynamically with time. For all experiments described below, we used a light intensity of 110 μW/cm^2^ for 16 h to control biofilm formation unless stated otherwise.

**Fig. 2.**
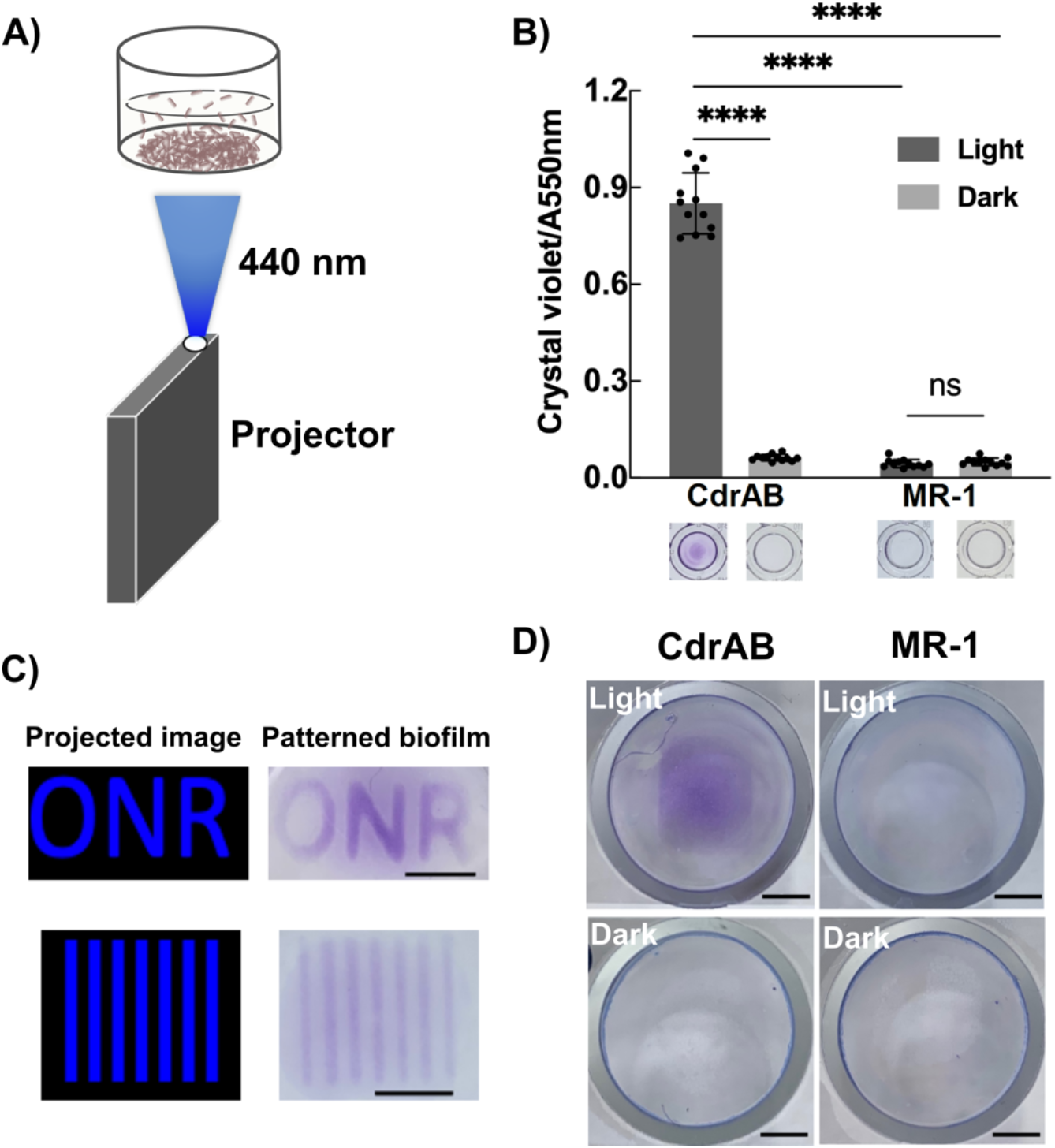
Light-induced biofilm formation and patterning. **A** Schematic for biofilm formation and patterning with blue light, performed with a portable LED projector. **B** Biofilm formation assay of CdrAB and wild type under blue light and dark conditions, respectively. n = 12 independent biological experiments per group. *p* < 0.0001 for CdrAB Light vs CdrAB Dark, CdrAB Light vs MR-1 Light and CdrAB Light vs MR-1 Dark (one-way ANOVA with Dunnett’s multiple comparisons test). *p* = 0.3466 for MR-1 Light vs MR-1 Dark (two-tailed unpaired *t* test). **C** Biofilm patterning of CdrAB in 12-well plates. **D** Biofilm patterning of CdrAB on ITO coated glass coverslip. Scale bars, 0.5 cm. Data are shown as mean ± SD. Significance is indicated as \*\*\*\**p* < 0.0001 and ns (not significant) *p* > 0.05.

Since CdrAB biofilm formation was significantly enhanced when grown under blue light illumination, we expected that the biofilm could also be patterned under blue light. We first cultured the cells in a 12-well polystyrene plate and illuminated various patterns on the bottom of the wells. Clear patterns (“ONR” and stripes) were visible after staining with crystal violet, which is used to identify regions of biofilm formation (Fig. 2C). We also optimized the illumination time (Supplementary Fig. 2 A and B) and growth medium volume (Supplementary Fig. 2 A and C) when patterning in the 12-well plates. Pattern formation was optimal when culturing in 1 mL defined minimal medium for 16 h, showing the sharpest contrast between exposed and unexposed regions of the plate, which was consistent with the results of biofilm formation assay described above (Supplementary Fig. 1B).

To test our patterning strategy on surfaces other than plastic, we moved to attempt biofilm patterning on glass surfaces by illuminating various patterns on the underside of glass bottom dishes. As shown in Supplementary Fig. 3, clear biofilm patterns were observed after crystal violet staining (Supplementary Fig. 3A). Microscopy revealed an increased and more uniform layer of cells in areas of the substrate exposed to blue light (Supplementary Fig. 3B). These results indicate the potential for patterning biofilms of *S. oneidensis* on a variety of substrates. For electrochemical measurements, biofilms would need to be patterned on electrodes. So next, we tested whether it was possible to pattern biofilms on transparent electrodes by culturing cells in a glass tube adhered to conductive, planar indium tin oxide (ITO) coated glass coverslip, a setup which can be used to make electrochemical measurements post-patterning. Since we observed that medium volume had an effect on biofilm patterning, we first optimized patterning in the glass tube setup by incubating different culture volumes in glass tubes adhered to cover glass with similar thickness to that of the ITO-coated coverslips. The patterned rectangular biofilms were most visible when 1 mL minimal medium was used to culture cells in the glass tube vessels (Supplementary Fig. 4). Then, we added 1 mL minimal medium to the glass tubes adhered to ITO coverslips and cultured the cells on the ITO surface by illuminating it with a rectangular pattern. As expected, the results showed that CdrAB biofilms can be patterned on the ITO surface under blue light illumination (Fig. 2D).

### The expression of CdrAB proteins does not significantly hinder extracellular electron transfer activity

Expressing outer membrane proteins on the cell surface of *S. oneidensis* may reduce the quantity of outer membrane cytochromes, which in turn may hinder extracellular electron transfer (EET)^43^. To verify whether the expression of CdrAB proteins would hinder the EET capability of *S. oneidensis*, we performed a ferrozine assay to compare the Fe^3+^ reduction capabilities of CdrAB and wild type strains after being cultured under blue light illumination and dark conditions, respectively. We also tested a mutant strain, *ΔMtr/ΔmtrB/ΔmtrE*, which lacks genes encoding periplasmic and outer-membrane cytochromes critical for EET, as a negative control^44^. Our results showed that the Fe^3+^ reduction of blue light illuminated CdrAB cells was initially slightly slower than that of CdrAB grown in the dark and wild type cultured either under blue light or in the dark, but then after a few hours, the extent of reduction was similar (Fig. 3A). This indicated that the expression of CdrAB proteins on the outer membrane of *S. oneidensis* may have interfered with the expression level or functionality of outer membrane cytochromes. A TMBZ heme staining SDS-PAGE gel was performed to determine the concentration of outer membrane cytochromes, which confirmed that CdrAB had a lower quantity of outer membrane cytochromes after being cultured under blue light (Supplementary Fig. 5). Overall, after expressing CdrAB proteins under blue light, the CdrAB strain maintained its EET capability and achieved a similar Fe^3+^ reduction level to that of the controls after 4 h (Fig. 3A).

**Fig. 3.**
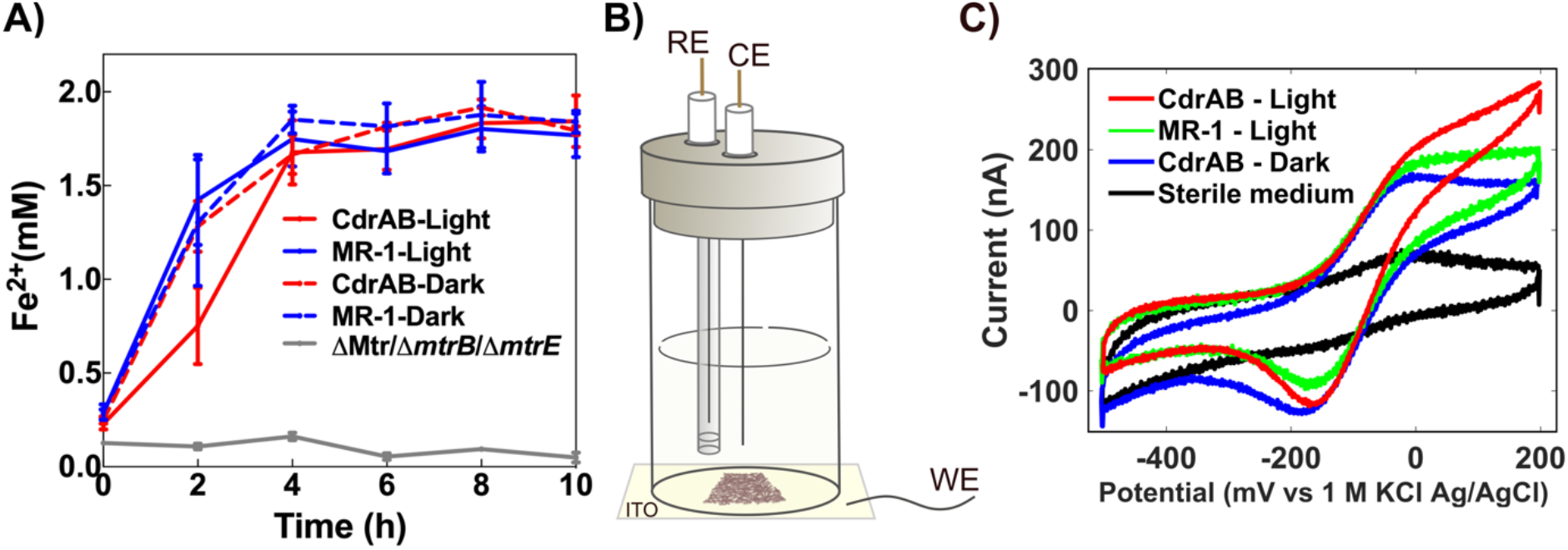
Extracellular electron transfer capability of CdrAB cells after being cultured under blue light and electrochemical activity for patterned CdrAB biofilms. **A** Ferrozine assay for CdrAB and wild type after being cultured under blue light and dark conditions, respectively. Data are shown as mean ± SD from 3 independent biological experiments. **B** Schematic of the bioreactor used for electrochemical measurements. **C** Representative cyclic voltammetry curves for CdrAB and wild type.

Since CdrAB cells could maintain EET activity after expressing CdrAB proteins, we reasoned that the patterned biofilms also remained electrochemically active. To verify this, CdrAB cells were incubated overnight in planar ITO-bottom glass tube reactors while a defined, rectangular light pattern was projected onto the base of the reactors from below (Figs. 2 A and D and 3B). As non-pattern-forming controls, glass tube reactors containing wild type cells exposed to the same projected light pattern and CdrAB cells kept in the dark were also incubated overnight. After overnight patterning and washing, the ITO bottom reactors were transferred to an anaerobic chamber for electrochemical measurements. From the cyclic voltammetry (CV) data, we observed that CdrAB, both under light and dark conditions, remained capable of electrode reduction at the same redox potential as the wild type (Fig. 3C). We also observed that, under the above experimental conditions, light patterned CdrAB produced more current than the dark condition and the wild type, likely due to the enhanced biofilm formation (Fig. 3C). Biofilm patterning was also attempted under anaerobic conditions to improve cytochrome expression. We observed that biofilm patterning was still viable under anaerobic conditions (Supplementary Fig. 6B) and that the CdrAB strain was again capable of electrode reduction at the same redox potential as the wild type (Supplementary Fig. 7). However, we observed increased cell attachment in the unexposed regions after anaerobic patterning (Supplementary Fig. 6). Thus, all subsequent light-patterning was performed under aerobic conditions prior to anaerobic electrochemical measurements.

### Electrochemical activity and conductance of patterned biofilms is proportional to biofilm surface area

With CdrAB capable of electrochemical activity when patterned onto ITO, we investigated electrochemical activity of patterned biofilms as a function of biofilm size. Three different sizes of biofilms were patterned onto custom ITO interdigitated array (IDA)-bottom glass tube reactors by projecting rectangular light patterns with the following dimensions: 1.3 x 0.25 cm^2^ (small pattern), 1.3 x 0.5 cm^2^ (medium pattern), and 1.3 x 1 cm^2^ (large pattern). We then used electrochemical gating to measure the biofilm conduction current for the three different pattern sizes. Here, the conduction current of patterned biofilms bridging the source and drain working electrodes (WEs) of a custom ITO IDA (Fig. 4A) was measured as a function of the potentials swept at each IDA WE. A fixed offset was maintained between the electrodes to serve as a driving force for conduction^17^. Crystal violet staining after the gating measurements clearly showed that three different sizes of biofilms were patterned (Fig. 4B). From the raw IDA source (*S*) and drain (*D*) CVs of our gating measurements, we observed increasing electrochemical activity as a function of increasing patterned biofilm size (Supplementary Fig. 8). By plotting the conduction current (*I_cond_*) vs the gate potential (*E_G_*, average potential of WEs during the gating scan), our results showed peak-shaped curves centered at the redox potential of *Shewanella* EET proteins (Fig. 4C), as expected for a redox conduction mechanism^17^, and indicating cell-to-cell long distance electron transport within light patterned biofilms. From our size dependent *I_cond_* vs *E_G_* data, we observed increasing conduction current as a function of increasing biofilm size (Fig. 4C), as expected since cells form the conduction channel between the source and drain electrodes. In essence, a larger biofilm forms a larger “wire” to carry the current between the source and drain. We also patterned the same sizes of biofilms onto planar ITO-bottom glass tube reactors ahead of CV measurements. These CVs gave results similar to those of our raw IDA source and drain CVs, showing increasing electrochemical activity as a function of increasing biofilm size, with the cells cultured in the dark giving the lowest current response (Supplementary Fig. 9). Due to the large working area of our custom IDA, compared to previously used commercial IDAs, and our ability to pattern continuous biofilms, compared to the patchy monolayers formed by wild type MR-1, we were able to detect tenfold larger conduction currents than what was previously measured in *S. oneidensis*^16^.

**Fig. 4.**
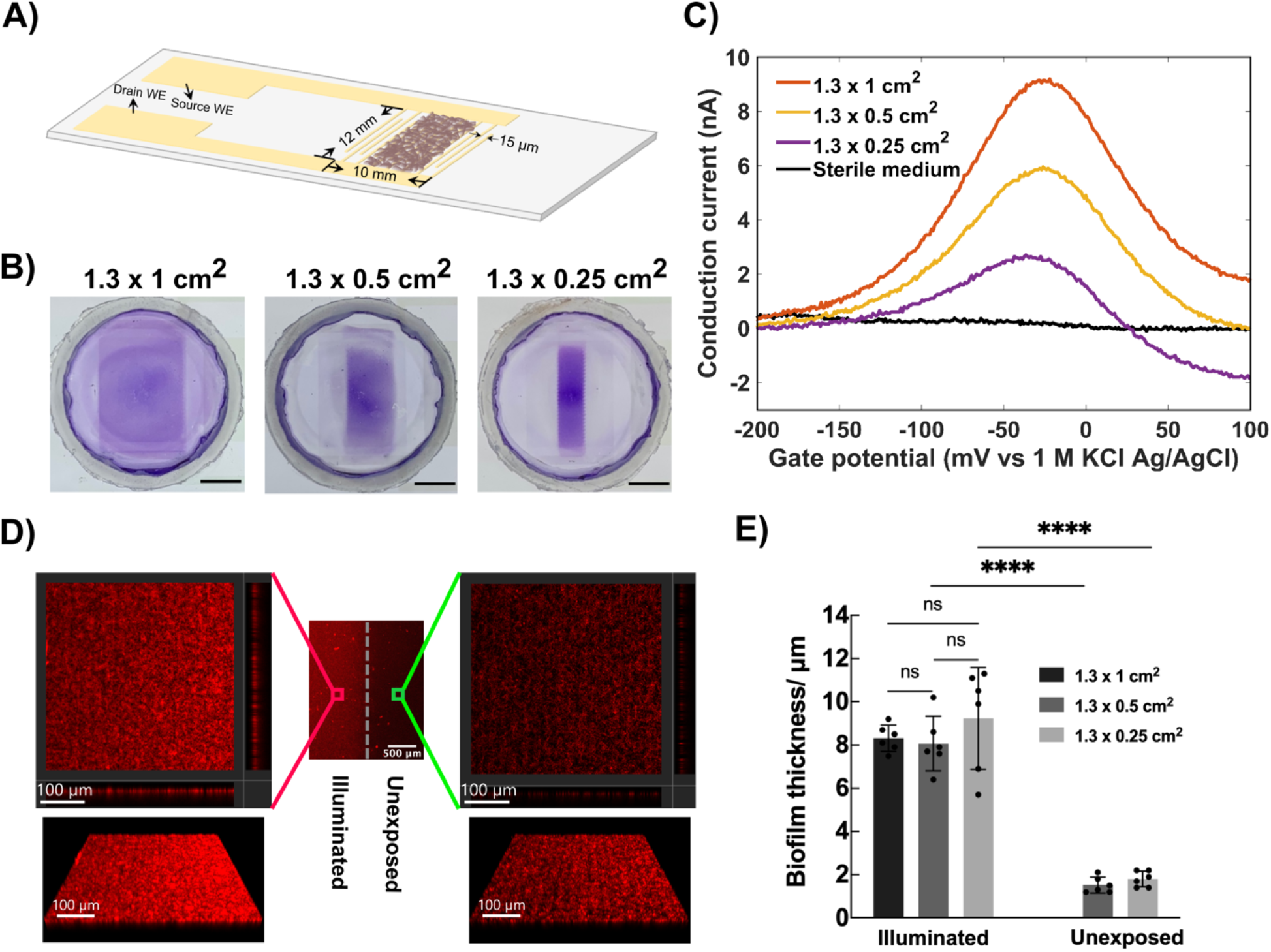
Electrochemical gating measurements for patterned CdrAB biofilms of different sizes. **A** Diagram of the custom transparent ITO IDA electrode. **B** Crystal violet staining of patterned biofilms with different sizes after the gating measurements, Scale bars, 0.5 cm. **C** Representative conduction currents curves of patterned biofilms with different sizes calculated from electrochemical gating measurements. **D** Confocal microscope images of biofilms on ITO IDA in blue-light illuminated region and unexposed region (1.3 x 0.25 cm^2^ pattern sample). **E** Thicknesses of biofilms on ITO IDAs in blue-light illuminated region and unexposed region of the patterned biofilm. n = 6 average biofilm thickness values calculated from 6 cross sections of three different confocal microscopy images for per group. The method for biofilm thickness analysis is shown in Supplementary methods and Supplementary Fig. 10. *p* = 0.9587 for illuminated 1.3 x 1 cm^2^ vs 1.3 x 0.5 cm^2^, *p* = 0.5862 for illuminated 1.3 x 0.1 cm^2^ vs 1.3 x 0.25 cm^2^ and *p* = 0.4291 for illuminated 1.3 x 0.5 cm^2^ vs 1.3 x 0.25 cm^2^ (one-way ANOVA with Dunnett’s multiple comparisons test). *p* < 0.0001 for 1.3 x 0.5 cm^2^ illuminated vs unexposed and 1.3 x 0.25 cm^2^ illuminated vs unexposed (two-tailed unpaired *t* test). Data are shown as mean ± SD. Significance is indicated as \*\*\*\**p* < 0.0001 and ns (not significant) *p* > 0.05.

### Defined pattern geometry enables biofilm conductivity to be calculated

After the electrochemical gating measurements, thicknesses of biofilms on ITO IDAs were obtained from confocal microscopy (Supplementary Fig. 10), which showed that a uniform and approximately 10 μm thick biofilm was formed in the patterned region for all three sizes used, with a thinner (~2 μm thick), sparse layer of cells present off the illuminated regions (Fig. 4 D and E). With a defined biofilm shape enabled by light-patterning, along with confocal microscopy biofilm thickness measurements and the conduction current measured via electrochemical gating, we were able to calculate a conductivity value for living CdrAB biofilms (see Supplementary methods). Using the dimensions and conduction current of the large biofilm pattern, a conductivity on the order of 4 nS/cm was calculated. The model used^45^ to calculate conductivity requires the conduction channel bridging the source and drain of an interdigitated electrode to be of uniform thickness. Thus, the large pattern size was used for this calculation, because a biofilm of uniform coverage and thickness was patterned over the entire interdigitated area in this case. For the small and medium patterned biofilms, only a portion of the interdigitated area was covered with the pattern, while the rest of the electrode area was covered with a thinner, patchy cell layer. This created a biofilm conduction channel of non-uniform thickness in the cases of the small and medium patterns.

## Discussion

We developed a strategy for patterning electroactive bacterial biofilms through the expression of CdrAB aggregation proteins under the control of the blue light-induced pDawn genetic circuit. In this work, several different aggregation proteins were tested with the pDawn genetic circuit since it was not clear which aggregation proteins could mediate adhesion of *S. oneidensis* cells. Of the aggregation proteins tested, only CdrAB proteins encouraged cell-cell adhesion of *S. oneidensis*. Previous studies have shown that CdrA is transported by CdrB to the cell surface and plays a common role as a biofilm matrix cross-linker with different extracellular exopolysaccharides (EPS) to promote aggregation of *P. aeruginosa* cells^35, 46^. Due to the close genetic relationship between *Pseudomonas* and *Shewanella*, CdrA may also interact with the EPS of *S. oneidensis* and enable cell aggregation. Although CdrAB expression enabled spatial control over biofilm formation, the cytochrome expression of *Shewanella* was decreased. This resulted in slightly lower EET activity, an issue that warrants further attention.

Bacterial biofilm patterning using optogenetic tools has been achieved with several approaches on different substrates without surface pretreatment at high spatial resolution^31, 32^. However, these patterned biofilms have not been naturally electrochemically active. While studies have been done to impart conductivity on *E. coli* patterned biofilms through interfacing self-assembly curli fibers with gold nanoparticles^47, 48^, such approaches require modifications of patterned biofilms with conductive nanoparticles. Our patterning strategy allows us to pattern naturally electroactive *S. oneidensis* biofilms without significantly hindering electrochemical capabilities of the strain.

Prior efforts to direct electroactive biofilm formation on surfaces have focused solely on engineering cell-electrode attachment, rather than patterning, through synthetic biology and materials engineering strategies. These works were based on either (I) enhancing biofilm formation by expressing adhesive appendages on cell surfaces^49, 50^ and increasing *c*-di-GMP levels^51, 52^ or (II) placing complementary chemical or DNA-based structures on substrates and cell surfaces to bond cells to electrodes^53–57^. Compared with these previous works, our strategy does not require electrode pretreatment for electroactive biofilm patterning and can generate robust biofilms with defined dimensions. As demonstrated in this work, this technique can enable tunable biofilm conduction and electrochemical activity by controlling the amount and location of electroactive cells on unmodified transparent working electrodes.

Importantly, precise control over biofilm geometry enabled us, for the first time, to quantitatively extract the intrinsic conductivity (~ 4 nS/cm) of living *Shewanella* biofilms. This was not possible previously as *Shewanella* typically makes patchy monolayers on electrode surfaces rather than forming uniform biofilms with defined dimensions, which are required for calculating conductivty^16^. With our controlled patterning strategy, it is now possible to compare *S. oneidensis* conductivity measurements to recent kinetic Monte Carlo simulations of a newly proposed collision-exchange electron transport mechanism^58^. In that work, micrometer-scale conduction in *S. oneidensis* was proposed to arise from the lateral diffusion of cytochromes, leading to collisions and inter-protein electron exchange along cell membranes. Utilizing experimentally determined outer membrane diffusion coefficients and electron hopping rates of *S. oneidensis* cytochromes, simulated conductivity was found to be on the order of 7 nS/cm, in excellent agreement with the conductivity value found in this work. If implemented in other exoelectrogens, our patterning technique could enable conductivity to be calculated for other electroactive biofilms and thus allow for more accurate modeling of biofilm conduction. Additionally, combining biofilm patterning with known biofilm conductivity can enable designer electrochemical activity and conduction in microbe-based bioelectronics as conductivity allows electron transport to be predicted for a given pattern size.

Although light-induced expression of CdrAB enabled the formation of dense biofilms in blue light-exposed regions of electrode surfaces, cells still attached to unexposed areas of the electrode and produced measurable currents. Future work and applications may benefit from improvements of the current-pattern resolution for defined biofilm dimensions by reducing attachment of cells in unexposed regions. This may also enable size-dependent current generation of anaerobically patterned biofilms, which are likely to be more conductive than aerobically grown biofilms due to their enhanced cytochrome expression. We have tried to remove the cells in the unexposed regions with orbital shaking to increase shear forces during the washing steps, but there remained cells in the unexposed regions. In future work, synthetic biology strategies could be developed to remove or kill nonspecifically surface-attached cells.

Our work provides an avenue for better understanding the relationship between electrochemical activity and electroactive biofilm geometry. Our work also opens the possibility for similar methods to be used to pattern electroactive biofilms of other exoelectrogens. Some approaches have been developed to enhance the EET of native electroactive biofilms by combining them with conductive nanoparticles^59, 60^. Thus, patterned biofilms could act as a scaffold for directed conductive nanoparticles biosynthesis or for nanoparticle-biofilm doping to further increase the EET capabilities of patterned biofilms for other applications. In short, we have developed a facile technique to direct the deposition of living conductive biofilms in relation to solid-state electrodes and we anticipate that this approach will enable further innovations for both studying and harnessing bioelectronics, akin to the role that traditional photolithography played in the development of solid-state electronics.

## Methods

### Bacterial strains, plasmids, and growth conditions

*Escherichia coli* DH5α was used for plasmids construction. *Pseudomonas aeruginosa* PAO1 was used as the source for *cdrAB* genes amplification. *Shewanella oneidensis* MR-1 was used as the host strain.

Plasmid pDawn-Ag43 which contains a blue-light-sensor system and expresses an *E. coli* aggregation protein Ag43 was obtained from *Addgene*^31^. Plasmids pDawn-mCherry, pDawn-AggA and pDawn-CdrAB were generated through replacing gene *ag43* with *mCherry, aggA* and *cdrAB*, respectively (Fig. 1A). Then the above four plasmids were transformed independently into *S. oneidensis* MR-1 via electroporation to generate the corresponding strains. All plasmids, strains and primers used in this study are listed in Supplementary Table 1.

### Fluorescence measurement of mCherry and cell clumping assay

Overnight cultures (1%, v/v) of mCherry strain were transferred into 5 mL fresh LB broth and grown to late log phase (OD_600_nm about 1-1.5) under the dark condition to prevent undesired photoactivation. Then, the cultures (1%, v/v) were seeded into 5 mL of LB broth and incubated at 30 °C either under the blue light or dark condition while shaking at 200 rpm. Blue light was provided by attaching LED strip lights to the wall inside the shaker. Cultures were collected after overnight incubation (16 h) to measure the fluorescence of mCherry and the cell optical density (OD_600_nm). Quantitative mCherry measurements were performed via a plate reader (Infinite 200 PRO, Tecan) at an excitation wavelength of 590 nm and an emission wavelength of 650 nm. The OD_600_nm was determined using a spectrophotometer (Spectronic 200, Thermo scientific). Relative fluorescence intensity was calculated by normalization against per OD_600_nm of whole cells. Fluorescence of mCherry was imaged via fluorescent microscope equipped with 100× oil immersion objective lens (Eclipse Ti, Nikon).

For the cell clumping assay, late log phase cultures (OD_600_nm about 1-1.5) of *Shewanella* strains containing different aggregation proteins were transferred into (1%, v/v) 5 mL fresh minimal medium and incubated overnight (16 h) at 30 °C under either blue light or dark condition while shaking at 200 rpm. Then, the cultures were left to rest for 30 mins. The upper region cultures of the tube were collected to measure the OD_top_. Then, the cultures were rigorously vortexed for 10 secs and the OD_total_ was determined. The aggregation index was defined as 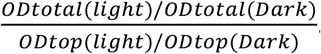.

### Biofilm formation assay and patterning on plastic and glass

The late log phase LB cultures (OD_600_nm about 1-1.5) were diluted into fresh minimal medium to an OD_600_nm of about 0.01. 100 μL of the dilution was added to each well in a 96-well plate for biofilm assays^42^. 1 mL and 3 mL of the dilution were added to each well of a 12-well plate and to the glass bottom dish for biofilm patterning, respectively. The vessels containing the cultures were taped to the ceiling of a 30 °C incubator. A portable smart projector (A5 Pro, Wowoto) was secured below the vessels in the incubator and pointed up at the bottom surface of the vessels (Fig. 2A). 440 nm blue light patterns created in Microsoft PowerPoint were projected onto the underside of the culture vessels.

After the overnight culturing of cells for different times and under different intensities of bluelight illumination, the medium was subsequently discarded. The patterned biofilms were gently washed three times with PBS. Then biofilms formed on the wells and dishes were stained with 0.1% crystal violet for 10 mins, rinsed three times with PBS and dried at room temperature for imaging. Before crystal violet staining, the light-induced biofilm patterned area and unexposed region in the same glass bottom dish were observed via light microscopy. Finally, 30% acetic acid was added to dissolve the absorbed crystal violet for 10 mins and A550 nm quantification of the solution was determined by plate reader.

An adjustable neutral density filter (KF01.063, K&F Concept) was used for tuning the illumination intensity by placing it at the aperture of the projector. An optical power meter (PM100USB, Thorlabs) was used for measuring the illumination intensity of the projected patterns.

For patterning on the surface of indium tin oxide (ITO) coated glass coverslips and custom ITO IDAs ahead of electrochemical measurements, 1 mL of diluted cells were added to the bioreactor, which was made by attaching a 20 mm diameter and 2.5 cm tall glass tube on an electrode. During the incubation, the glass tubes were sealed with a microporous membrane filter (AirOtop Seals). After 16 h of patterning, medium was discarded, and the patterned biofilms were washed five times, for 10 mins each time, with PBS or non-turnover minimal medium in a shaker at 220 rpm to reduce the non-specific patterned cells.

### Resting cell ferrozine assay

Resting cell ferrozine assay was modified based on a previous study^22^. Cells from late log phase LB cultures were diluted into 5 mL fresh minimal medium to an OD_600_nm of about 0.01. Then, cells were incubated for 16 h at 30 ^o^C under either blue light or dark condition while shaking at 200 rpm. Cells were collected by centrifuging at 4,200 rpm, 4 ^o^C for 15 mins and then washed with fresh minimal medium once. After that, cells were inoculated into sealed serum bottles containing 25 mL anaerobic minimal medium to an OD_600_nm of about 0.1. 2 mM ferric citrate was added into the anaerobic minimal medium as the electron acceptor. The samples were incubated at 30 ^o^C without shaking. Every two hours, 10 μL of each sample was added immediately to 90 μL 1 M HCl in a 96-well plate followed by 100 μL 0.01% ferrozine. Then, after mixing the samples well and letting them sit for 10 mins, the absorbances of the samples at 562 nm was determined with a plate reader. A standard curve of freshly made ferrous sulfate was used to determine the Fe^2+^ concentrations.

### Transparent-bottom bioreactor construction

To observe the surface coverage and thickness of patterned biofilms on bioreactor working electrodes (WEs) *in situ*, transparent-bottom bioreactors were constructed. Planar, 22 mm by 40 mm commercial indium tin oxide (ITO) coated glass coverslips (Prod no. 06494-AB, Structure Probe, Inc.) or custom ITO IDAs (designed in house and fabricated via foundry service) were used as the bioreactor base. Before use, the commercial ITO glass coverslips and the custom IDAs were rinsed with acetone, isopropanol, and then with DI water, and then dried with N_2_. Thin copper wires (Prod no. 1227, TCS 20’ 32 gauge wire) were electrically connected to the WEs with silver paint (Prod no. 16035, TED PELLA, Inc.). To mechanically strengthen the wire-electrode connections, they were covered with epoxy (Gorilla Glue Co.). Functioning as the body of the bioreactors, glass tubes (2.5 cm tall with a 19 mm and 22 mm inner and outer diameter, respectively) were adhered overtop of the WEs with siliconized sealant (DAP Kwik Seal Ultra Premium Siliconized Sealant) (Fig. 3B). Custom, PEEK plastic lids were used with the transparent-bottom reactors along with custom Pt wire counter electrodes (CEs) and 1 M KCl Ag/AgCl reference electrodes (REs) (CHI111P, CH Instruments, Inc.) (Fig. 3B). All potentials reported in this manuscript are vs 1 M KCl Ag/AgCl.

### Measurements of electrochemical activity and biofilm conduction

All electrochemical measurements were performed in an anaerobic chamber (Bactron 300, Sheldon Manufacturing, Inc.) with a 95:5 (N_2_/H_2_) atmosphere. Sterile medium (blank) electrochemical measurements were performed before reactors were used in biofilm patterning. For the commercial ITO reactors, minimal media was used for the blank measurements, while non-turnover (NT) minimal media was used for the interdigitated array (IDA) blank measurements. During the patterning steps, the CEs and REs were removed from the reactors and the reactors were simply used as a culturing vessel. For the commercial ITO reactors, blank and biofilm cyclic voltammetry (CV) measurements were performed from −500 mV to either 200 or 300 mV at 1 mV/sec using a four channel Squidstat (Admiral Instruments). To perform the electrochemical gating done with the IDA reactors, two Gamry Reference 600 potentiostats connected with a synchronization cable and operating in bipotentiostat mode were used. Blank gating scans were also performed before patterning. For the blank and biofilm gating measurements, the IDA working electrodes (WEs) were scanned at rate of 1 mV/sec from −500 mV to 300 mV but with a fixed gating offset *V_SD_*= 20 mV, where *V_SD_* = *E_D_* – *E_S_* and where *E_D_* and *E_S_* are the potentials at each of the IDA WEs. To determine *I_cond_*, we first assumed that equal and opposite sign *I_cond_* plus background currents of the same sign were measured at each IDA WE. Then, *I_cond_* could be calculated by subtracting the drain and source currents and dividing by two, (*I_D_* – *I_S_*)/2. Three cycles were performed for both the CV and gating scans and only data from the third cycle is presented in this manuscript. Ahead of anaerobic patterning, the reactors were sealed with sterile rubber stoppers before being removed from the anaerobic chamber. After aerobic patterning, inside the anaerobic chamber, the reactor media was exchanged for anaerobic media before all electrochemical measurements. Prior to the size-dependent CV and gating scans, chronoamperometry (CA) was performed on the reactors for 1 to 6 h with the WEs held at 200 mV.

### Statistical analysis

All statistical analyses were performed by the Prism software (version 9.0; GraphPad) using one-way ANOVA with Dunnett’s multiple comparisons test or two-tailed unpaired *t* test. All data are presented as the mean ± SD. *P* values in all graphs were generated with tests as indicated in figure legends and are represented as follows: \**p* < 0.05, \*\**p* < 0.01, \*\*\**p* < 0.001, \*\*\*\**p* < 0.0001 and ns (not significant) *p* > 0.05. For the representative experiments shown in figures, three independent experiments were repeated with similar results.

## Data availability

All relevant data supporting the key findings of this study are available within the article and Supplementary Information. Plasmids and strains generated for this study will be shared upon request to the corresponding author.

## Acknowledgments

We thank the Nano3 Cleanroom at the University of California San Diego for enabling fabrication of the electrodes. We thank Karla Abuyen for the help of modifying illustrations and Cesar Rodriguez for preliminary work on this project. This study was supported by the US Office of Naval Research Multidisciplinary University Research Initiative Grant No. N00014-18-1-2632.

## Author Contributions

F.Z., M.S.C., J.A.G., M.Y.E.-N. and J.Q.B. designed research; F.Z., M.S.C., K.L.N. and C.M.C. performed research; F.Z., M.S.C. and K.L.N. analyzed data; and F.Z., M.S.C., M.Y.E.-N. and J.Q.B. wrote the paper. All authors edited the manuscript.

## References

1. Myers, C. R. & Nealson, K. H. Bacterial manganese reduction and growth with manganese oxide as the sole electron acceptor. Science 240, 1319–1321 (1988).

2. Lovley, D. R., Stolz, J. F., Nord, G. L. & Phillips, E. J. P. Anaerobic production of magnetite by a dissimilatory iron-reducing microorganism. Nature 330, 252–254 (1987).

3. Shi, L. et al. Extracellular electron transfer mechanisms between microorganisms and minerals. Nat. Rev. Microbiol. 14, 651–662 (2016).

4. McMillan, D. G. G., Marritt, S. J., Butt, J. N.& Jeuken, L. J. C. Menaquinone-7 is specific cofactor in tetraheme quinol dehydrogenase CymA. J. Biol. Chem. 287, 14215–14225 (2012).

5. Edwards, M. J. et al. Structural modeling of an outer membrane electron conduit from a metal-reducing bacterium suggests electron transfer via periplasmic redox partners. J. Biol. Chem. 293, 8103–8112 (2018).

6. Fonseca, B. M. et al. Mind the gap: Cytochrome interactions reveal electron pathways across the periplasm of *Shewanella oneidensis* MR-1. Biochem. J. 449, 101–108 (2013).

7. Hartshorne, R. S. et al. Characterization of an electron conduit between bacteria and the extracellular environment. Proc. Natl. Acad. Sci. U. S. A. 106, 22169–22174 (2009).

8. White, G. F. et al. Rapid electron exchange between surface-exposed bacterial cytochromes and Fe(III) minerals. Proc. Natl. Acad. Sci. U. S. A. 110, 6346–6351 (2013).

9. Edwards, M. J., White, G. F., Butt, J. N., Richardson, D. J. & Clarke, T. A. The crystal structure of a biological insulated transmembrane molecular wire. Cell 181, 665–673.e10 (2020).

10. Marsili, E. et al. *Shewanella* secretes flavins that mediate extracellular electron transfer. Proc. Natl. Acad. Sci. U. S. A. 105, 3968–3973 (2008).

11. Coursolle, D., Baron, D. B., Bond, D. R. & Gralnick, J. A. The Mtr respiratory pathway is essential for reducing flavins and electrodes in *Shewanella oneidensis*. J. Bacteriol. 192, 467–474 (2010).

12. Pirbadian, S. et al. *Shewanella oneidensis* MR-1 nanowires are outer membrane and periplasmic extensions of the extracellular electron transport components. Proc. Natl. Acad. Sci. U. S. A. 111, 12883–12888 (2014).

13. Xu, S., Jangir, Y. & El-Naggar, M. Y. Disentangling the roles of free and cytochrome-bound flavins in extracellular electron transport from *Shewanella oneidensis* MR-1. Electrochim. Acta 198, 49–55 (2016).

14. Zacharoff, L. A. & El-Naggar, M. Y. Redox conduction in biofilms: From respiration to living electronics. Curr. Opin. Electrochem. 4, 182–189 (2017).

15. Snider, R. M., Strycharz-Glaven, S. M., Tsoi, S. D., Erickson, J. S. & Tender, L. M. Long-range electron transport in *Geobacter sulfurreducens* biofilms is redox gradient-driven. Proc. Natl. Acad. Sci. U. S. A. 109, 15467–15472 (2012).

16. Xu, S., Barrozo, A., Tender, L. M., Krylov, A. I. & El-Naggar, M. Y. Multiheme cytochrome mediated redox conduction through *Shewanella oneidensis* MR-1 cells. J. Am. Chem. Soc. 140, 10085–10089 (2018).

17. Yates, M. D. et al. Thermally activated long range electron transport in living biofilms. Phys. Chem. Chem. Phys. 17, 32564–32570 (2015).

18. Logan, B. E. & Rabaey, K. Conversion of wastes into bioelectricity and chemicals by using microbial electrochemical technologies. Science 337, 686–690 (2013).

19. Tanzil, A. H. et al. Production of gold nanoparticles by electrode-respiring *Geobacter sulfurreducens* biofilms. Enzyme Microb. Technol. 95, 69–75 (2016).

20. Zhang, Y., Hsu, L. H. H. & Jiang, X. Living electronics. Nano Res. 13, 1205–1213 (2020).

21. Yim, S. S. et al. Robust direct digital-to-biological data storage in living cells. Nat. Chem. Biol. 17, 246–253 (2021).

22. West, E. A., Jain, A. & Gralnick, J. A. Engineering a native inducible expression system in *Shewanella oneidensis* to control extracellular electron transfer. ACS Synth. Biol. 6, 1627–1634 (2017).

23. Cao, Y., Li, X., Li, F. & Song, H. CRISPRi-sRNA: Transcriptional-translational regulation of extracellular electron transfer in *Shewanella oneidensis*. ACS Synth. Biol. 6, 1679–1690 (2017).

24. Li, F. H. et al. Developing a population-state decision system for intelligently reprogramming extracellular electron transfer in *Shewanella oneidensis*. Proc. Natl. Acad. Sci. U. S. A. 117, 23001–23010 (2020).

25. Jensen, H. M. et al. Engineering of a synthetic electron conduit in living cells. Proc. Natl. Acad. Sci. U. S. A. 107, 19213–19218 (2010).

26. Jensen, H. M., TerAvest, M. A., Kokish, M. G., & Ajo-Franklin, C. M. CymA and exogenous flavins improve extracellular electron transfer and couple it to cell growth in Mtr-expressing *Escherichia coli*. ACS Synth. Biol. 5, 679–688 (2016).

27. Su, L. et al. Modifying cytochrome c maturation can increase the bioelectronic performance of engineered *Escherichia coli*. ACS Synth. Biol. 9, 115–124 (2020).

28. Levskaya, A. et al. Engineering *Escherichia coli* to see ligh. Nature 438, 441–442 (2005).

29. Ohlendorf, R., Vidavski, R. R., Eldar, A., Moffat, K. & Möglich, A. From dusk till dawn: One-plasmid systems for light-regulated gene expression. J. Mol. Biol. 416, 534–542 (2012).

30. Tabor, J. J., Levskaya, A. & Voigt, C. A. Multichromatic control of gene expression in *Escherichia coli*. J. Mol. Biol. 405, 315–324 (2011).

31. Jin, X. & Riedel-Kruse, I. H. Biofilm Lithography enables high-resolution cell patterning via optogenetic adhesin expression. Proc. Natl. Acad. Sci. U. S. A. 115, 3698–3703 (2018).

32. Moser, F., Tham, E., González, L. M., Lu, T. K. & Voigt, C. A. Light-controlled, high-resolution patterning of living engineered bacteria onto textiles, ceramics, and plastic. Adv. Funct. Mater. 29, 1–11 (2019).

33. Huang, Y., Xia, A., Yang, G. & Jin, F. Bioprinting living biofilms through optogenetic manipulation. ACS Synth. Biol. 7, 1195–1200 (2018).

34. Chen, F. & Wegner, S. V. Blue-light-switchable bacterial cell-cell adhesions enable the control of multicellular bacterial communities. ACS Synth. Biol. 9, 1169–1180 (2020).

35. Borlee, B. R. et al. *Pseudomonas aeruginosa* uses a cyclic-di-GMP-regulated adhesin to reinforce the biofilm extracellular matrix. Mol. Microbiol. 75, 827–842 (2010).

36. Reichhardt, C., Wong, C., da Silva, D. P., Wozniak, D. J. & Parsek, M. R. CdrA interactions within the *Pseudomonas aeruginosa* biofilm matrix safeguard it from proteolysis and promote cellular packing. MBio 9, 1–12 (2018)

37. Pu, L., Yang, S., Xia, A. & Jin, F. Optogenetics manipulation enables prevention of biofilm formation of engineered *Pseudomonas aeruginosa* on surfaces. ACS Synth. Biol. 7, 200–208 (2018).

38. Henderson, I. R., Meehan, M. & Owen, P. Antigen 43, a phase-variable bipartate outer membrane protein, determines colony morphology and autoaggregation in *Escherichia coli* K-12. FEMS Microbiol. Lett. 149, 115–120 (1997).

39. Danese, P. N., Pratt, L. A., Dove, S. L. & Kolter, R. The outer membrane protein, Antigen 43, mediates cell-to-cell interactions within *Escherichia coli* biofilms. Mol. Microbiol. 37, 424–432 (2000).

40. De Vriendt, K. et al. Proteomics of *Shewanella oneidensis* MR-1 biofilm reveals differentially expressed proteins, including AggA and RibB. Proteomics 5, 1308–1316 (2005).

41. De Windt, W. et al. AggA is required for aggregation and increased biofilm formation of a hyper-aggregating mutant of *Shewanella oneidensis* MR-1. Microbiology 152, 721–729 (2006).

42. O’Toole, G. A. Microtiter dish Biofilm formation assay. J. Vis. Exp., 10–11 (2010).

43. Kane, A. L., Bond, D. R. & Gralnick, J. A. Electrochemical analysis of *Shewanella oneidensis* engineered to bind gold electrodes. ACS Synth. Biol. 2, 93–101 (2013).

44. Coursolle, D. & Gralnick, J. A. Reconstruction of extracellular respiratory pathways for iron(III) reduction in *Shewanella oneidensis* strain MR-1. Front. Microbiol. 3, 1–11 (2012).

45. Kankare, J. & Kupila, E. L. In-situ conductance measurement during electropolymerization. J. Electroanal. Chem. 322, 167–181 (1992).

46. Reichhardt, C. et al. The versatile *Pseudomonas aeruginosa* biofilm matrix protein CdrA promotes aggregation through different extracellular exopolysaccharide interactions. J. Bacteriol. 202, 1–9 (2020).

47. Chen, A. Y. et al. Synthesis and patterning of tunable multiscale materials with engineered cells. Nat. Mater. 13, 515–523 (2014).

48. Wang, X. et al. Programming cells for dynamic assembly of inorganic nano-objects with spatiotemporal control. Adv. Mater. 30, 1–10 (2018).

49. Liu, X. et al. Flagella act as *Geobacter* biofilm scaffolds to stabilize biofilm and facilitate extracellular electron transfer. Biosens. Bioelectron. 146, 111748 (2019).

50. Leang, C., Malvankar, N. S., Franks, A. E., Nevin, K. P. & Lovley, D. R. Engineering *Geobacter sulfurreducens* to produce a highly cohesive conductive matrix with enhanced capacity for current production. Energy Environ. Sci. 6, 1901–1908 (2013).

51. Liu, T. et al. Enhanced *Shewanella* biofilm promotes bioelectricity generation. Biotechnol. Bioeng. 112, 2051–2059 (2015).

52. Hu, Y., Wu, Y., Mukherjee, M. & Cao, B. A near-infrared light responsive *c*-di-GMP module-based AND logic gate in *Shewanella oneidensis*. Chem. Commun. 53, 1646–1648 (2017).

53. Catania, C., Karbelkar, A. A. & Furst, A. L. Engineering the interface between electroactive bacteria and electrodes. Joule 5, 743–747 (2021).

54. Furst, A. L., Smith, M. J., Lee, M. C. & Francis, M. B. DNA hybridization to interface current-producing cells with electrode surfaces. ACS Cent. Sci. 4, 880–884 (2018).

55. Suo, D., Fang, Z., Yu, Y. Y. & Yong, Y. C. Synthetic curli enables efficient microbial electrocatalysis with stainless-steel electrode. AIChE J. 66, 9–12 (2020).

56. Young, T. D. et al. Selective promotion of adhesion of *Shewanella oneidensis* on mannose-decorated glycopolymer surfaces. ACS Appl. Mater. Interfaces 12, 35767–35781 (2020).

57. Lienemann, M. et al. Towards patterned bioelectronics: facilitated immobilization of exoelectrogenic *Escherichia coli* with heterologous pili. Microb. Biotechnol. 11, 1184–1194 (2018).

58. Chong, G. W. et al. Single molecule tracking of bacterial cell surface cytochromes reveals dynamics that impact long-distance electron transport. bioRxiv. DOI: 10.1101/2021.11.02.466829 (2021).

59. Jiang, X. et al. Nanoparticle facilitated extracellular electron transfer in microbial fuel cells. Nano Lett. 14, 6737–6742 (2014).

60. Yang, C. et al. Carbon dots-fed *Shewanella oneidensis* MR-1 for bioelectricity enhancement. Nat. Commun. 11, 1–11 (2020).

