## Supplementary Information for "Light-induced Patterning of Electroactive Bacterial Biofilms"

**Supplementary methods**

**Growth conditions**

*E. coli* DH5α was cultivated in Luria-Bertani (LB) medium (10 g/L tryptone, 5 g/L yeast extract and 5 g/L NaCl) at 37 ^o^C, 200 rpm. *S. oneidensis* strains were cultivated in LB medium or minimal medium^1^ (15.1 g/L PIPES buffer, 3.4 g/L NaOH, 1.5 g/L NH_4_Cl, 0.1 g/L KCl, 0.6 g/L NaH_2_PO_4_·H_2_O, 3.36 g/L 60% (w/w) NaC_3_H_5_O_3_, 1 mL mineral solution, 1 mL vitamin solution^2^, 10 mL amino acid solution, pH 7.0) at 30 °C, 200 rpm. The mineral solution consists of 20 g/L C_6_H_9_NO_3_ (dissolved with NaOH to pH 8.0), 30 g/L MgSO_4_·7H_2_O, 5 g/L MnSO_4_·H_2_O, 10 g/L NaCl, 1 g/L FeSO_4_·7H_2_O, 1 g/L CaCl_2_·2H_2_O, 1 g/L CoCl_2_·6H_2_O, 1.3 g/L ZnCl_2_, 0.1 g/L CuSO_4_·5H_2_O, 0.1 g/L AlK(SO_4_)_2_·12H_2_O, 0.1 g/L H_3_BO_3_, 0.25 g/L Na_2_MoO_4_·2H_2_O, 0.24 g/L NiCl_2_·6H_2_O and 0.25 g/L Na_2_WO_4_·2H_2_O. The vitamin solution consists of 0.02 g/L C_10_H_16_N_2_O_3_S, 0.02 g/L C_19_H_19_N_7_O_6_, 0.1 g/L C_8_H_12_ClNO_3_, 0.05 g/L C_17_H_20_N_4_O_6_, 0.05 g/L C_18_H_18_Cl_2_N_4_OS, 0.05 g/L C_6_H_5_NO_2_, 0.05 g/L C_9_H_16_NO_5_·1/2Ca, 0.001 g/L C_63_H_88_CoN_14_O_14_P, 0.05 g/L C_7_H_7_NO_2_, and 0.05 g/L C_8_H_14_O_2_S_2_. The amino acid solution consists of 2 g/L L-glutamic acid, 2 g/L L-arginine, and 2 g/L DL-Serine. When necessary, media were supplemented with spectinomycin (Spec, 100 μg/mL).

When MR-1 strains were cultured under anaerobic conditions, Na_2_C_4_H_2_O_4_ was added to the above minimal media. The medium used in this paper for electrochemical measurements is prepared using the above minimal medium recipe without vitamin solution. When performing electrochemical gating measurements, non-turnover (NT) minimal medium is used without vitamin solution, NaC_3_H_5_O_3_, and Na_2_C_4_H_2_O_4_. The media used all for electrochemical measurements were purged with nitrogen.

**Tetramethylbenzidine (TMBZ) heme stain SDS-PAGE protein gel**

The late log phase LB cultures (OD_600nm_ about 1-1.5) of *S. oneidensis* strains were transferred into (1%, v/v) 5 mL fresh minimal medium and incubated overnight (16 h) at 30 ^o^C under either blue light or dark condition while shaking at 200 rpm. Then, the cells of 1 mL culture with an OD_600nm_ of 2.0 were collected. The cell pellet was resuspended to 150 μL SDS sample buffer without β-mercaptoethanol. Subsequently, the sample was boiled for 10 mins and centrifuged with 13,000 rpm for 5 mins. 20 μL of sample was loaded for running a 12% SDS-PAGE protein gel. The gel was rinsed with water for 5 mins and then immersed in a solution which contained 15 mL 6.3 mM TMBZ and 35 mL 0.25 M sodium acetate (pH 5.0). After keeping the sample in the dark for 2 h, 3 mL 30% hydrogen peroxide was added and staining was visible within 3 minutes.

**Microfabrication of custom interdigitated array electrodes**

The custom indium tin oxide (ITO) interdigitated array (IDA) electrodes were designed in house and were fabricated by the University of California, Sand Diego Nano3 cleanroom foundry service using standard photolithography techniques. The interdigitated area of the devices consisted of 200 pairs of electrodes fingers, with 12 mm long and 10 mm wide fingers and a 15 μm gap between fingers. Thus, each device had a total working electrode area of 0.48 cm^2^. The fabrication process is briefly outlined as follows. 100 mm diameter and 500 μm thick BK-7 glass wafers (University Wafer, Inc.) were used as the electrode substrate. Solvent cleaned wafers were coated with photoresist (PR) and then a laser writer was used to project three IDA electrode patterns, side by side, onto the PR with UV light. PR developer was then used to remove the light exposed PR and then 300 nm of ITO was sputter coated onto the wafers. Solvents were used to remove the excess ITO and PR, leaving just the electrode pattern. To improve the conductivity of the electrodes, the wafers were baked in an N_2_ furnace at 400 ^o^C for one hour^3^. As a final step, the wafers were coated in PR and then diced into three 24 mm by 60 mm chips, with each chip containing one ITO IDA pattern. It must also be noted that since the interdigitated area of these devices (including fingers and gaps) was 1.2 x 1 cm^2^, the effective biofilm pattern sizes used in our electrochemical gating measurement are 1 mm smaller lengthwise than the projected patterns.

**Additional technical notes about the electrochemical gating measurements.**

Electrochemical gating is often performed under non-turnover (NT) conditions to remove the catalytic contributions to the source and drain currents. Before the size-dependent gating experiments were done, an additional experiment was performed in an IDA reactor, with a biofilm pattern, to confirm that the patterned biofilms were under NT conditions during all gating measurements. Blank gating, patterning, and reactor media exchanges were performed as described earlier. For this experiment, the large CdrAB biofilm pattern size was used. This was done to ensure that an upper limit could be determined for the time it would take all patterned biofilms to achieve NT conditions. When the biofilm patterned IDA was brought into the anaerobic chamber, an initial gating scan was performed and then a chronoamperometry (CA) scan was performed for 3 days, with the working electrodes (WEs) held at 200 mV. Gating scans were then periodically performed on the reactor, approximately every 6-18 h to monitor the shape of the cyclic voltammetry (CV) curves over time. As biofilm CV scans change from “S” shaped to duck shaped, this mechanistically signifies that electron transport between the biofilm and the WEs is no longer governed by metabolic activity, but rather by redox cycling of the cell surface-bound cytochromes and the WEs^4^. After the three days, the shape of the CV curves from the gating scans did not change, indicating the culture began under non-turnover conditions. Thus, before each gating scan, only a 1 to 6 hr NT CA was done. Additionally, when calculating the biofilm conductivity using *I_cond_*, a linear relationship between *I_cond_* and *V_SD_* is assumed. To confirm this, for the medium biofilm pattern size, in addition to the 20 mV offset gating scan, a 10 mV and 30 mV offset gating scan were performed. Then, the peak *I_cond_* for the three different gating offset scans were plotted against the *V_SD_* used for each scan to generate a *I_cond_* vs *V_SD_* plot, which was observed to be linear (Supplementary Fig. 11).

***In situ* microscopy observations of patterned biofilms on electrodes**

After electrochemical measurements, to stain the cells, the reactor media was discarded and 2 mL of minimal medium containing FM4-64X membrane dye was added to the bioreactors and left to sit for 15 mins. Then the biofilms on the electrode were imaged using either the 40× or 100× objective lens of a Nikon Eclipse Ti inverted fluorescent microscope. To prepare confocal microscopy samples of patterned biofilms on the IDA electrodes, FM4-64X dye containing minimal medium was added to the reactors and left to sit for 30 mins. Subsequently, the medium was exchanged for 3% glutaraldehyde to fix the cells at 4 ^o^C overnight. The samples were then washed with PBS for 3 times before imaging. Confocal microscopy was performed using a Zeiss LSM 880 inverted microscope equipped with an argon laser and a 20× dry objective lens. 3D image analysis was performed using the software Imaris.

**Biofilm thickness analysis**

In order to determine the biofilm thickness of patterned biofilms, cross-sectional confocal images were processed and analyzed. First, two cross sections in the plane of the biofilm were taken from each confocal microscopy image. Next, a threshold was applied to cross-sectional confocal images to convert them to binary images. The threshold was determined by examining pixels in the dark background of each confocal microscopy image, far away from the biofilm itself. The biofilm thickness at a particular slice along the binary cross-section was reported by summing over non-zero pixels at that slice. By summing over all slices, the average biofilm thickness of that cross section can be determined. Three confocal microscopy images were analyzed for each sample.

**Calculation of patterned biofilm conductivity**

After confirming the linear relationship between *I_cond_* and *V_SD_* (Supplementary Fig. 11) for our patterned biofilms, Ohm’s Law (equation 1) was used to determine biofilm conductance (*G*). Where *G* is equal to the intrinsic conductivity of the biofilm (*σ*) and a geometric factor (*S*) dependent on the electrode and biofilm geometries (equation 2). Here *I_cond_* is given by the peak conduction current for the large-sized pattern, with baseline correction, and *V_SD_* is equal to the 20 mV potential offset used during the electrochemical gating measurements. Adapting a conformal mapping method, equation 3 was used to approximate *S* for an IDA consisting of 200 pairs of fingers with 10 μm finger widths and with 15 μm finger gaps^5^. The geometric factor used was for the case where the biofilm thickness was on the order of the size of the IDA finger gaps. In equation 3, *l* is equal to the length of the gap that snakes between the interdigitated fingers and can be calculated by multiplying the number of gaps (399 gaps) by the length of each gap (1.2 cm). The term “*a*” is equal to half the width of a finger gap and the term “*g*” is equal to the biofilm thickness. With the conductance determined from the electrochemical gating results and the geometric factor determined by the IDA geometry and biofilm thickness, equations 2 and 3 are used to solve for the biofilm conductivity.

$I_{cond}=GV_{SD}$	(1) $G=\sigma S$	(2)

$S=\frac{l}{\pi}ln(\frac{8g}{\pi a})$	(3)

**Supplementary figures**

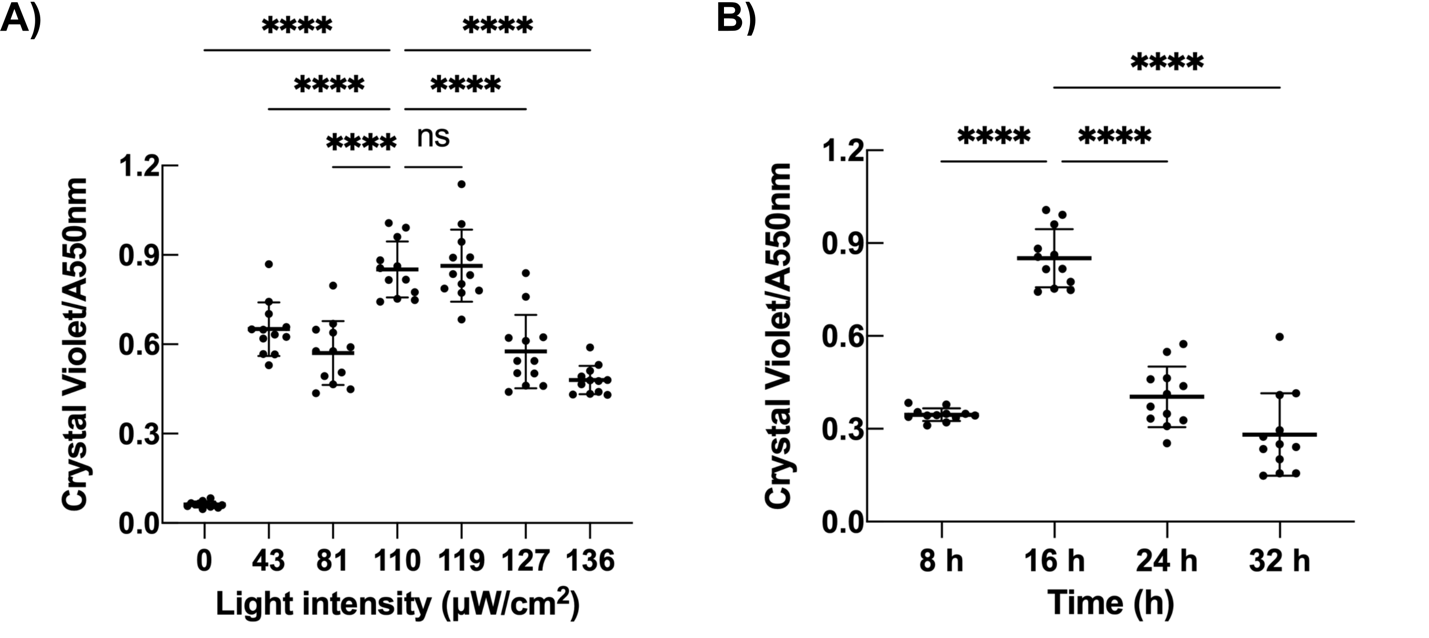

**Supplementary Fig. 1 Optimization of the biofilm formation under different light intensities and illumination time**. **A** Biofilm formation under different light intensities. n=12 independent biological experiments per group. *p* = 0.9982 for 110 μW/cm^2^ vs 119 μW/cm^2^. *p* < 0.0001 for 110 μW/cm^2^ vs all the remaining groups. **B** Biofilm formation with different illumination time. n = 12 independent biological experiments per group. *p* < 0.0001 for 16 h vs 8 h, 16 h vs 24 h and 16 h vs 32 h. Data are shown as mean ± SD. Statistical analysis was performed using one-way ANOVA with Dunnett’s multiple comparisons test and significance is indicated as *****p* < 0.0001 and ns (not significant) *p* > 0.05.

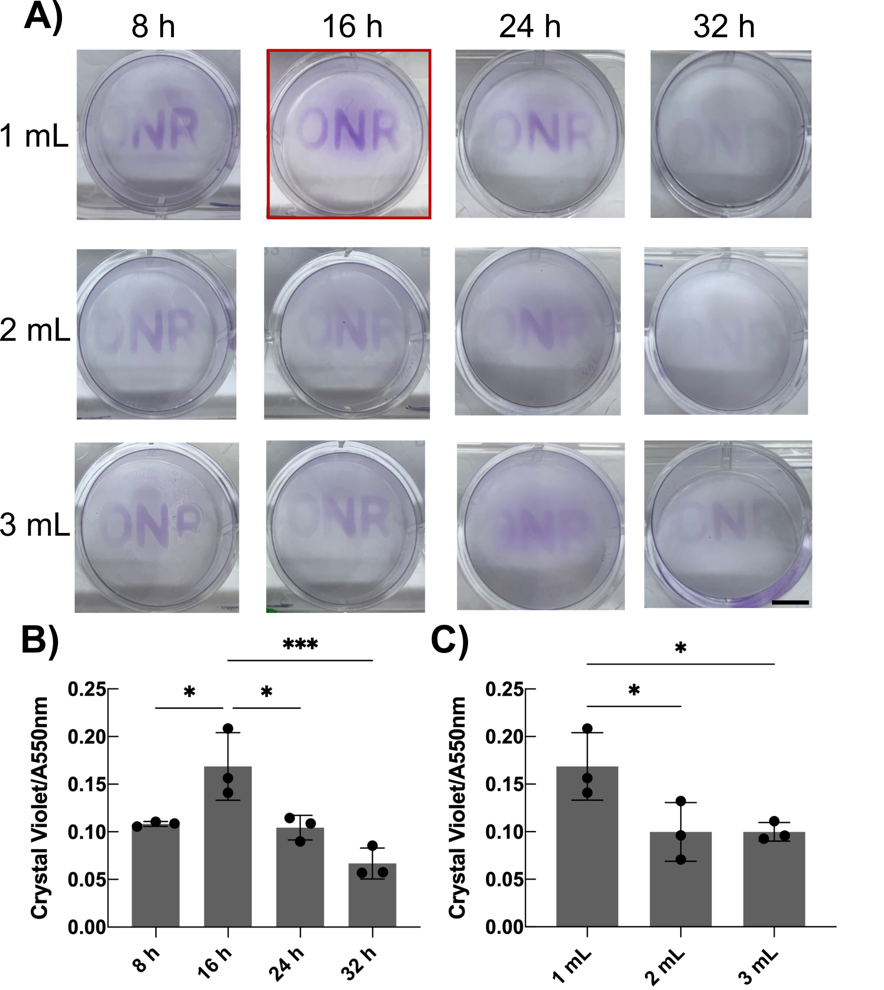

**Supplementary Fig. 2 Biofilm patterning on plastic substrates**. **A** Optimization of biofilm patterning in 12-well plates. Scale bar 0.5 cm. **B** Crystal violet measurement of biofilm patterning in 12-well plates with different time. n = 3 independent biological experiments per group. *p* = 0.0175 for 16 h vs 8 h, *p* = 0.0127 for 16 h vs 24 h and *p* = 0.0008 for 16 h vs 32 h. **C** Crystal violet measurement of biofilm patterning in 12-well plates with different culture volumes. n = 3 independent biological experiments per group. *p* = 0.0395 for 1 mL vs 2 mL and *p* = 0.0398 for 1 mL vs 3 mL. Data are shown as mean ± SD. Statistical analysis was performed using one-way ANOVA with Dunnett’s multiple comparisons test and significance is indicated as **p* < 0.05 and ****p* < 0.001.

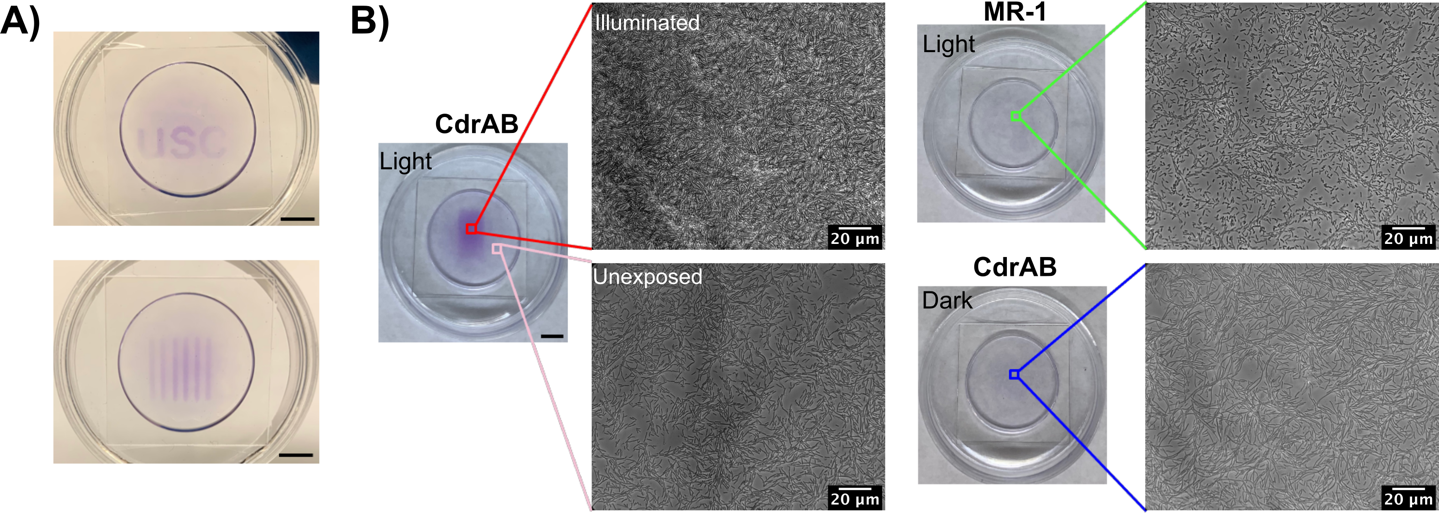

**Supplementary Fig. 3 Biofilm patterning on glass substrates**. **A** Patterned biofilms (“usc” and stripes) on glass bottom dishes. **B** Microscopy images of biofilms on the glass bottom dishes for CdrAB and wild type strains illuminated with the same rectangle and CdrAB strain cultured under the dark condition. Scale bars, 0.5 cm for the crystal violet stained patterned biofilm images.

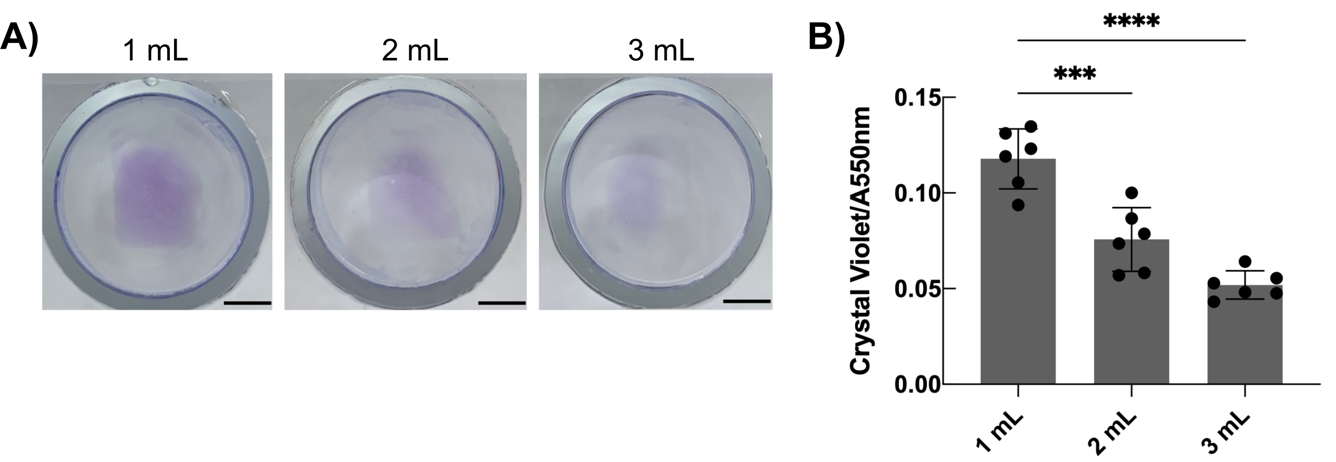

**Supplementary Fig. 4 Optimization of biofilm patterning using glass tubes**. **A** and **B** Crystal violet staining and measurement of biofilm patterning using glass tubes attached to cover glasses with different culture volumes. n = 6 independent biological experiments per group. *p* = 0.0002 for 1 mL vs 2 mL and *p* < 0.0001 for 1 mL vs 3 mL. Scale bars, 0.5 cm. Data are shown as mean ± SD. Statistical analysis was performed using one-way ANOVA with Dunnett’s multiple comparisons test and significance is indicated as ****p* < 0.001 and *****p* < 0.0001.

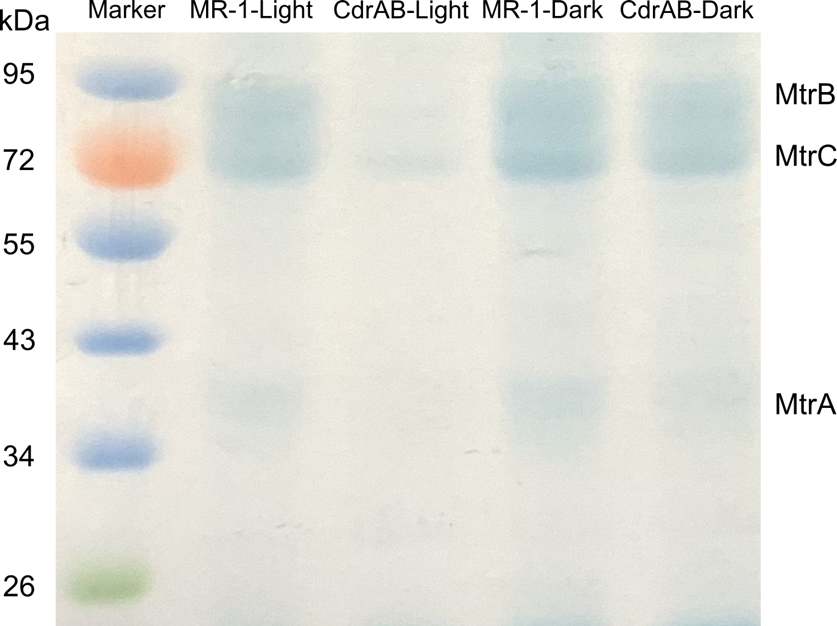

**Supplementary Fig. 5** TMBZ heme stain SDS-PAGE protein gel for CdrAB and wild type strains after being cultured aerobically under blue light illumination and dark conditions, respectively.

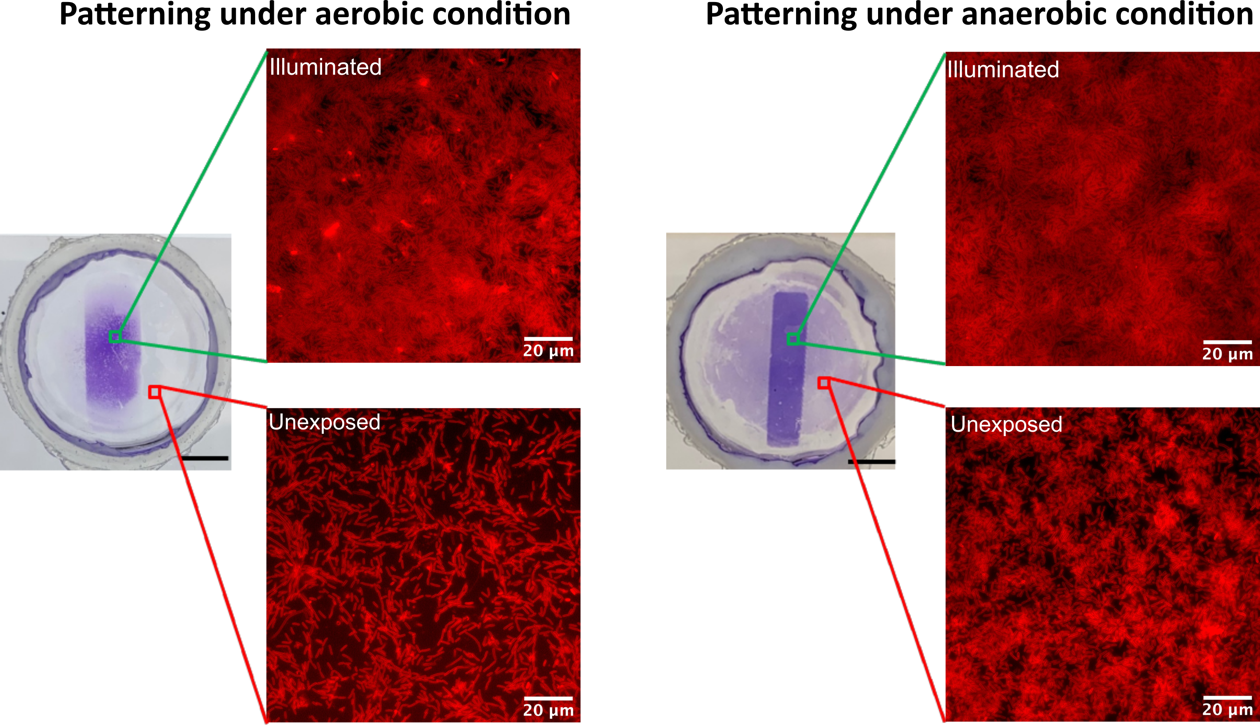

**Supplementary Fig. 6** Biofilm patterning on the ITO coated glass electrodes under aerobic and anaerobic conditions, respectively. Scale bars, 0.5 cm for the crystal violet stained patterned biofilm images. The cell attachment in the unexposed regions of the anaerobically patterned reactors was significantly larger than that observed in the aerobically patterned reactors.

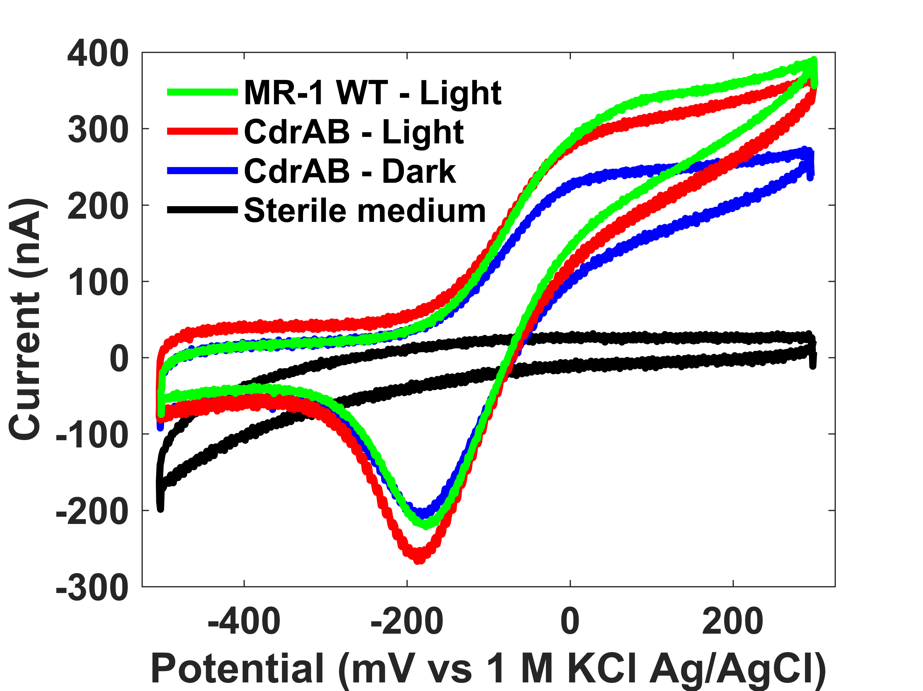

**Supplementary Fig. 7** Representative cyclic voltammetry curves for anaerobically patterned CdrAB and the anaerobically incubated, non-patterned wild type and CdrAB dark condition on ITO coated glass coverslips. While we observed that the light patterned CdrAB produces more current than the dark CdrAB condition, the WT produces more current than both the other conditions.

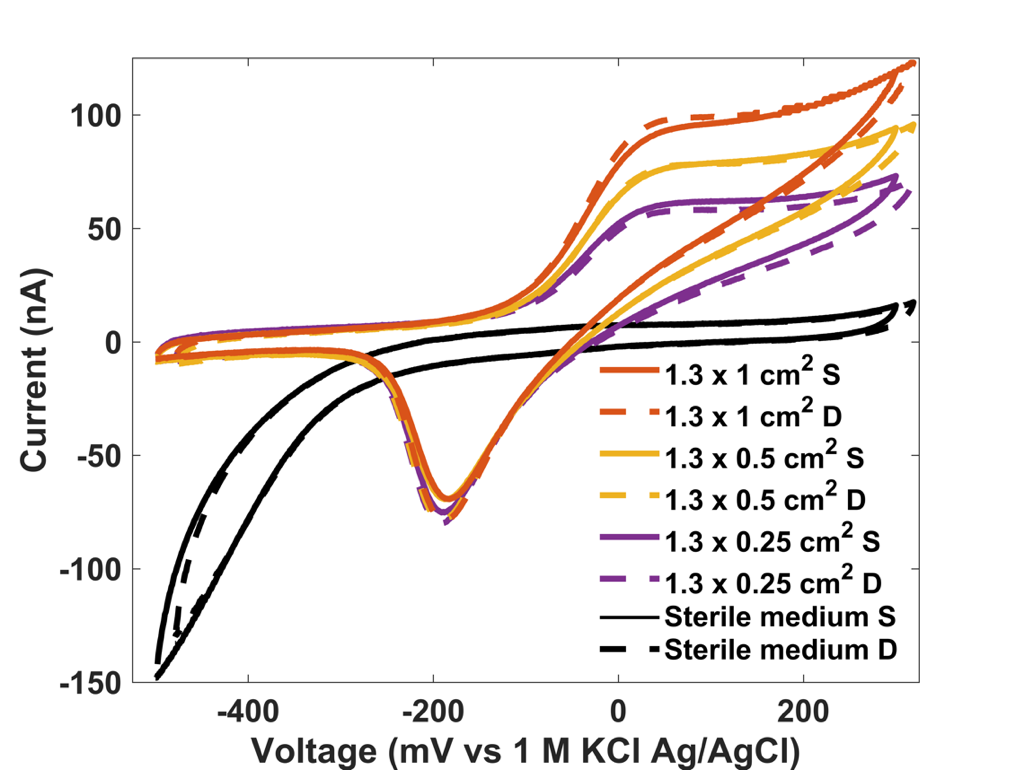

**Supplementary Fig. 8** Representative source and drain cyclic voltammetry curves for patterned CdrAB biofilms of different sizes on custom transparent ITO IDA electrodes. *S*: Source electrode. *D*: Drain electrode.

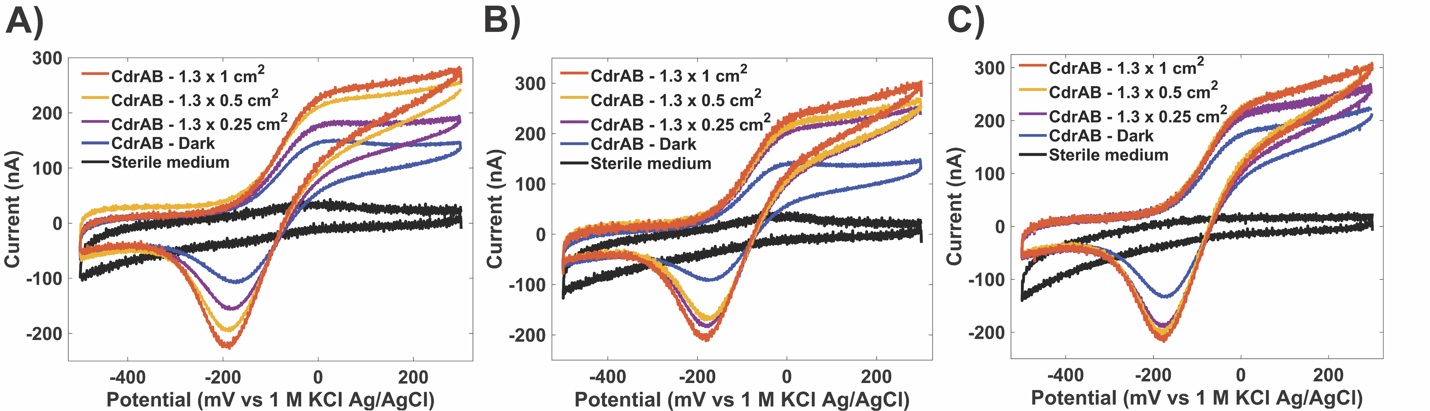

**Supplementary Fig. 9** **Triplicate cyclic voltammetry curves for patterned biofilms of CdrAB with different sizes on planar ITO electrodes.** In each replicate, the large pattern size (red) produces more current than the small pattern size (purple), which in turn produces more current than the non-patterned dark condition (blue). However, in **A**, the middle pattern size (orange) is shown to produce current between the values produced by the small and larger pattern. And then in **B** and in **C**, the current produced by the middle pattern size is shown to fluctuate between the values produced by the small and large patterns, respectively. This demonstrates the initial, non-optimized, biofilm size-dependent current resolution of our patterning technique.

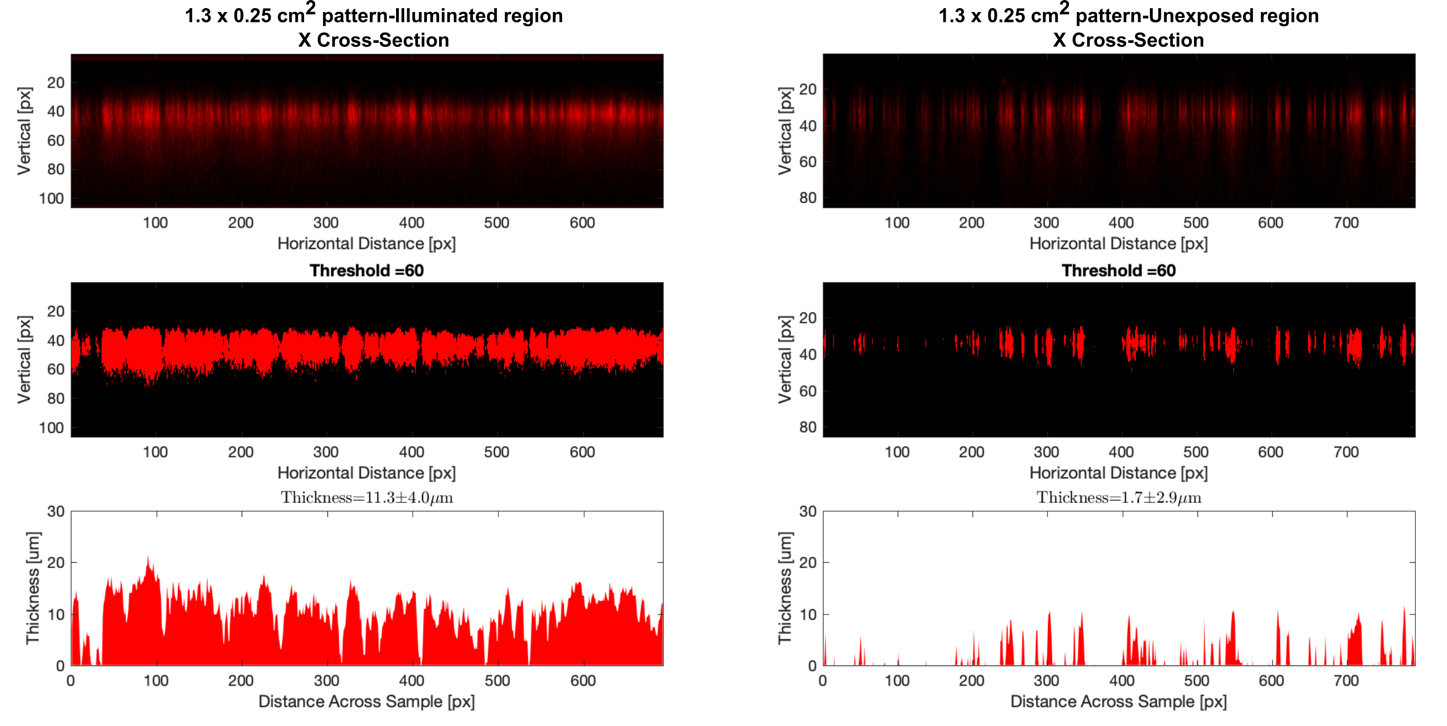

**Supplementary Fig. 10** **Examples for biofilm thickness measurements of illuminated regions and unexposed regions (1.3 x 0.25 cm^2^ pattern sample).**

*Top image*, the cross-sectional confocal image of a biofilm. The “vertical” axis is in the same direction as the biofilm, i.e. away from the substrate. The “horizontal” axis is in the plane of the biofilm, i.e. along the surface of the substrate.

*Middle image,* the top image with threshold applied. The number of red pixels along each horizontal slice is added and this determines the biofilm thickness.

*Bottom image,* the result of adding non-zero pixels after thresholding along the horizontal direction. The mean and standard deviation is displayed.

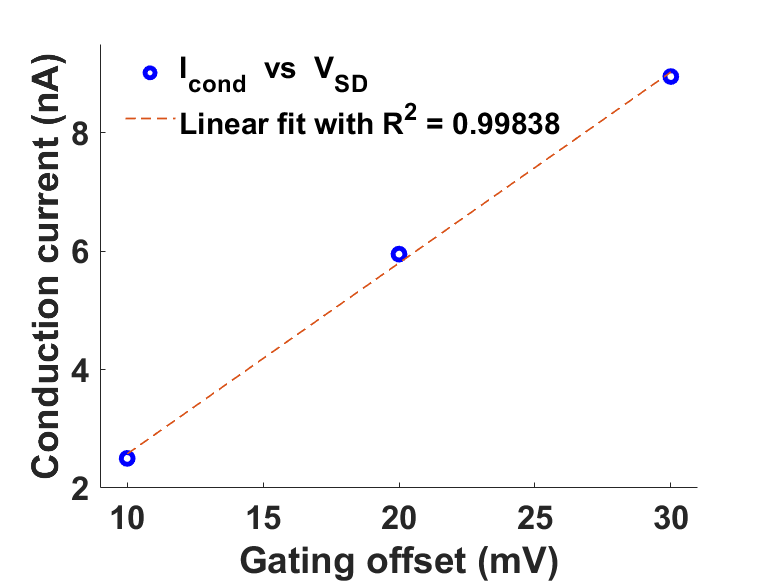

**Supplementary Fig. 11** Peak *I_cond_* vs varying gating offsets for the medium-sized (1.3 x 0.5 cm^2^) CdrAB pattern.

**Supplementary Table 1 Strains, plasmids and primers used in this study**

| **Strains, plasmids and primers** | **Description** | **Source or literature** |
| --- | --- | --- |
| **Strains** | | |
| MR-1 | *Shewanella oneidensis* MR-1 wild type strain | This work |
| mCherry | MR-1 derivative carrying pDawn-mCherry plasmid | This work |
| CdrAB | MR-1 derivative carrying pDawn-CdrAB plasmid | This work |
| Ag43 | MR-1 derivative carrying pDawn-Ag43 plasmid | This work |
| AggA  ΔMtr/Δ*mtrB*/Δ*mtrE* | MR-1 derivative carrying pDawn-AggA plasmid  MR-1 derivative without genes encoding periplasmic and outer membrane cytochromes critical for extracellular electron transfer | This work  (6) |
| **Plasmids** | | |
| pDawn-mCherry | Expression of mCherry under pDawn genetic circuit | This work |
| pDawn-CdrAB | Expression of CdrAB under pDawn genetic circuit | This work |
| pDawn-Ag43 | Expression of Ag43 under pDawn genetic circuit | (7) |
| pDawn-AggA | Expression of AggA under pDawn genetic circuit | This work |
| **Primers** | | |
| pDawn-mCherry-F | tcctcgcctttactcaccatTCTAGTAGGTTTCCTGTGTGAGG | This work |
| pDawn-mCherry-R | tggatgagctgtataaataaCTCGAGCACCACCACCAC | This work |
| mCherry-pDawn-F | tggtggtggtggtgctcgagTTATTTATACAGCTCATCCATAC | This work |
| mCherry-pDawn-R | cacacaggaaacctactagaATGGTGAGTAAAGGCGAG | This work |
| pDawn-CdrAB-F | atccggacggaccatTCTAGTAGGTTTCCTGTGTGAGG | This work |
| pDawn-CdrAB-R | gtggcgcgcttctgaCTCGAGCACCACCACCAC | This work |
| CdrAB-pDawn-F | gtggtggtgctcgagTCAGAAGCGCGCCACCAC | This work |
| CdrAB-pDawn-R | aggaaacctactagaATGGTCCGTCCGGATAGC | This work |
| pDawn-AggA-F | tactaaagtcttcatTCTAGTAGGTTTCCTGTGTGAGG | This work |
| pDawn-AggA-R | ggagcaaatcaatgaCTCGAGCACCACCACCAC | This work |
| AggA-pDawn-F | gtggtggtgctcgagTCATTGATTTGCTCCTCC | This work |
| AggA-pDawn-R | aggaaacctactagaATGAAGACTTTAGTAAGACG | This work |

**Supplementary references**

1.	Bretschger, O. et al*.* Current production and metal oxide reduction by *Shewanella oneidensis* MR-1 wild type and mutants. *Appl. Environ. Microbiol.* **73**, 7003–7012 (2007).

2.	Kieft, T. L. et al*.* Dissimilatory reduction of Fe(III) and other electron acceptors by a *Thermus* isolate. *Appl. Environ. Microbiol.* **65**, 1214–1221 (1999).

3.	Gross, B. J. & El-naggar, M. Y. A combined electrochemical and optical trapping platform for measuring single cell respiration rates at electrode interfaces. *Rev. Sci. Instrum.* **86**, 064301–1 to 064301–8 (2015).

4.	Xu, S., Barrozo, A., Tender, L. M., Krylov, A. I. & El-Naggar, M. Y. Multiheme Cytochrome Mediated Redox Conduction through *Shewanella oneidensis* MR-1 Cells. *J. Am. Chem. Soc.* **140**, 10085–10089 (2018).

5.	Kankare, J. & Kupila, E. L. In-situ conductance measurement during electropolymerization. *J. Electroanal. Chem.* **322**, 167–181 (1992).

6.	Coursolle, D. & Gralnick, J. A. Reconstruction of extracellular respiratory pathways for iron(III) reduction in *Shewanella oneidensis* strain MR-1. *Front. Microbiol.* **3**, 1–11 (2012).

7.	Jin, X. & Riedel-Kruse, I. H. Biofilm Lithography enables high-resolution cell patterning via optogenetic adhesin expression. *Proc. Natl. Acad. Sci. U. S. A.* **115**, 3698–3703 (2018).
